# Instant Effects of Semantic Information on Visual Perception

**DOI:** 10.1101/2023.04.19.537469

**Authors:** Alexander Enge, Franziska Süß, Rasha Abdel Rahman

**Author notes:** Correspondence concerning this article should be addressed to Alexander Enge, Rudower Chaussee 18, 12489 Berlin, Germany.

## Abstract

Does our perception of an object change once we discover what function it serves? We showed human participants (*n* = 48, 31 female, 17 male) pictures of unfamiliar objects either together with keywords matching their function, leading to semantically informed perception, or together with non-matching keywords, resulting in uninformed perception. We measured event-related potentials (ERPs) to investigate at which stages in the visual processing hierarchy these two types of object perception differed from one another. We found that semantically informed as compared to uninformed perception was associated with larger amplitudes in the N170 component (150–200 ms), reduced amplitudes in the N400 component (400–700 ms), and a late decrease in alpha/beta band power. When the same objects were presented once more without any information, the N400 and event-related power effects persisted, and we also observed enlarged amplitudes in the P1 component (100–150 ms) in response to objects for which semantically informed perception had taken place. Consistent with previous work, this suggests that obtaining semantic information about previously unfamiliar objects alters aspects of their lower-level visual perception (P1 component), higher-level visual perception (N170 component), and semantic processing (N400 component, event-related power). Our study is the first to show that such effects occur instantly after semantic information has been provided for the first time, without requiring extensive learning.

**Significance Statement:** There has been a long-standing debate about whether or not higher-level cognitive capacities such as semantic knowledge can influence lower-level perceptual processing in a top-down fashion. Here we could show for the first time that information about the function of previously unfamiliar objects immediately influences cortical processing within less than 200 ms. Of note, this influence does not require training or experience with the objects and related semantic information. Therefore, our study is the first to show effects of cognition on perception while ruling out the possibility that prior knowledge merely acts by pre-activating or altering stored visual representations. Instead, this knowledge seems to alter perception online, thus providing a compelling case against the impenetrability of perception by cognition.

## Introduction

Does our perception of an object change once we discover what function it serves? This question speaks to the long-standing debate about the cognitive (im)penetrability of perception by higher-level capacities such as semantic knowledge or language. According to one view, these cognitive capacities kick in only after the retinal input has been processed by a specialized module for visual perception (Fodor, 1983). This module is supposed to be encapsulated from higher-level inputs and processes visual information in a feed-forward fashion, progressing from lower to higher areas representing increasingly complex shapes and, eventually, whole objects (DiCarlo et al., 2012). This cognitive impenetrability hypothesis is challenged by another view, namely predictive coding theories that posit that higher-level areas influence ongoing perceptual processing early on by sending predictions down to lower-level areas (Churchland et al., 1994; Ahissar and Hochstein, 2004; Yuille and Kersten, 2006; Friston and Kiebel, 2009; Clark, 2013; Lupyan, 2015; Thierry, 2016; Teufel and Nanay, 2017; Lupyan et al., 2020). This view is supported by differences in ratings, detection rates, or reaction times for visual stimuli depending on their emotional (e.g., Phelps et al., 2016), linguistic (e.g., Slivac et al., 2021) or semantic (e.g., Gauthier et al., 2003) content. However, some of these studies received legitimate criticism, e.g., because comparisons between critical conditions included the confound of additionally comparing different visual stimuli, or for not being able to distinguish between perceptual or post-perceptual loci of tentative top-down effects based on behavioral measures (Firestone and Scholl, 2016).

Here we employ event-related potentials (ERPs) measured from the human EEG to mitigate most of these concerns: The temporal resolution of ERPs makes it possible to probe how early influences of high-level (e.g., semantic) information can be detected (Athanasopoulos and Casaponsa, 2020). When comparing objects after learning different amounts of semantic information about them, previous studies by our lab and others revealed differences in late ERP components associated with semantic processing (N400 component) and also in the visual P1 component (Abdel Rahman and Sommer, 2008; Samaha et al., 2018; Maier and Abdel Rahman, 2019; Weller et al., 2019). The early peak of the P1 and its source in the occipital cortex (Abdel Rahman and Sommer, 2008) point to an early effect of semantic knowledge on visual perception. It is less clear, however, if this effect acts in an online fashion (i.e., directly modulating perceptual processing, in accordance with predictive coding theories), or more indirectly, by altering stored visual object representations over the course of learning, which would then get reactivated once the object is reencountered later on (i.e., reflecting offline differences in visual object representations or prototypes that have been built up over the course of learning; Palmeri and Tarr, 2008).

To answer this question, we measured ERPs in response to unfamiliar objects directly while participants gained a semantic understanding of their function. We presented half of the objects with matching keywords, allowing participants to understand what kind of object they were viewing, and the other half with non-matching keywords, preventing participants from understanding what kind of object they were viewing. We then presented the same objects again to test for downstream effects of semantic information. We examined the influence of semantic information on ERPs associated with lower-level visual perception (P1 component), higher-level visual perception (N170 component), and semantic processing (N400 component). We hypothesized that semantic information would have instant effects on visual perception, by which we mean (a) that these effects can be observed on the same trial as the semantic information is being presented for the first time, and (b) that these effects are found not only in later, higher-level cognition-related ERP components (N400), but also in earlier, perception-related ERP components (P1 and/or N170), which would speak for predictive coding theories. We furthermore conducted an exploratory time-frequency analysis to test for effects on event-related power which, unlike ERP effects, do not need to be tightly phase-locked to stimulus onset.

## Materials and Methods

### Participants

Participants were 48 German native speakers (31 female, 17 male) with a mean age of 23.5 years (range 18–32 years) and no history of psychological disorder or treatment. No a priori power analysis was carried out. All participants were right-handed according to the Edinburgh inventory (Oldfield, 1971) and reported normal or corrected-to-normal vision. They provided written informed consent before starting the experiment and received a monetary compensation of €8 per hour for participating.

### Stimuli

Stimuli consisted of 240 grayscale photographs of real-world objects. Of these, 120 stimuli were well-known everyday objects (e.g., a bicycle, a toothbrush). These served as filler stimuli of no interest. The other 120 stimuli were rare objects presumed to be unfamiliar to most participants (e.g., a galvanometer, an udu drum; see Online Table 1 at https://doi.org/10.17605/osf.io/uksbc). All stimuli were presented on a light blue background with a size of 207 × 207 pixels on a 19-inch LCD monitor with a resolution of 1,280 × 1,024 pixels and a refresh rate of 75 Hz. At a standardized viewing distance of 90 cm, the images subtended approximately 3.9 degrees of participants’ horizontal and vertical visual angle. For each unfamiliar object, we created a pair of German keywords (a noun and a verb), describing the typical function or use of the object in a way that could be related to its visual features and their configuration (e.g., Stromstärke, messen [electric current, measuring]; Tonpott, trommeln [clay pot, drumming]; see Online Table 1). As our central experimental manipulation, half of the objects were presented together with keywords that matched their respective function, whereas the other half of the objects were presented together with non-matching keywords (which would have matched a different object). The matching keywords were expected to induce semantically informed perception, that is, participants suddenly understanding what kind of object they were viewing. The non-matching keywords were expected to prevent such an understanding and keep the perception of the object semantically uninformed. All participants saw each unfamiliar object with only one type of keywords (matching or non-matching). This assignment of keywords to objects was counterbalanced across participants so that each object was presented with matching keywords (leading to semantically informed perception) and non-matching keywords (leading to uninformed perception) to an equal number of participants. The experiment was programmed and displayed using Presentation® software (Neurobehavioral Systems, Inc., Berkeley, CA, www.neurobs.com).

As a manipulation check, we ran an online rating study where we presented 10 German speakers (3 female, 7 male, mean age = 25.3 years, range 20–35 years; none took part in the main study) with all 240 visual objects in random order and asked them to generate their own keywords that would describe the presumed function of the object. We used latent semantic analysis (Günther et al., 2019) with a word2vec embedding (deepset GmbH, Berlin, Germany, www.deepset.ai/german-word-embeddings) pretrained on the German Wikipedia to estimate semantic distances between these participant-generated keywords and the keywords used in our main EEG experiment (see Online Figures 1 and 2). In brief, it was substantially easier for participants to come with correct descriptions of familiar objects (mean cosine distance ± standard error = 0.78 ± 0.01) than for unfamiliar objects (0.66 ± 0.01), *t*(13) = 7.21, *p*_corr_ < .001 (linear mixed-effects model). In fact, this similarity between participant-generated keywords for the unfamiliar objects and the object-matching keywords presented in the main experiment was only slightly higher than the similarity between participant-generated keywords and the object-non-matching keywords presented in the main experiment (0.64 ± 0.01), *t*(24) = 5.09, *p*_corr_ < .001 (linear mixed-effects model), indicating that it was difficult for participants to know or guess the correct function of the unfamiliar objects.

**Figure 1.**
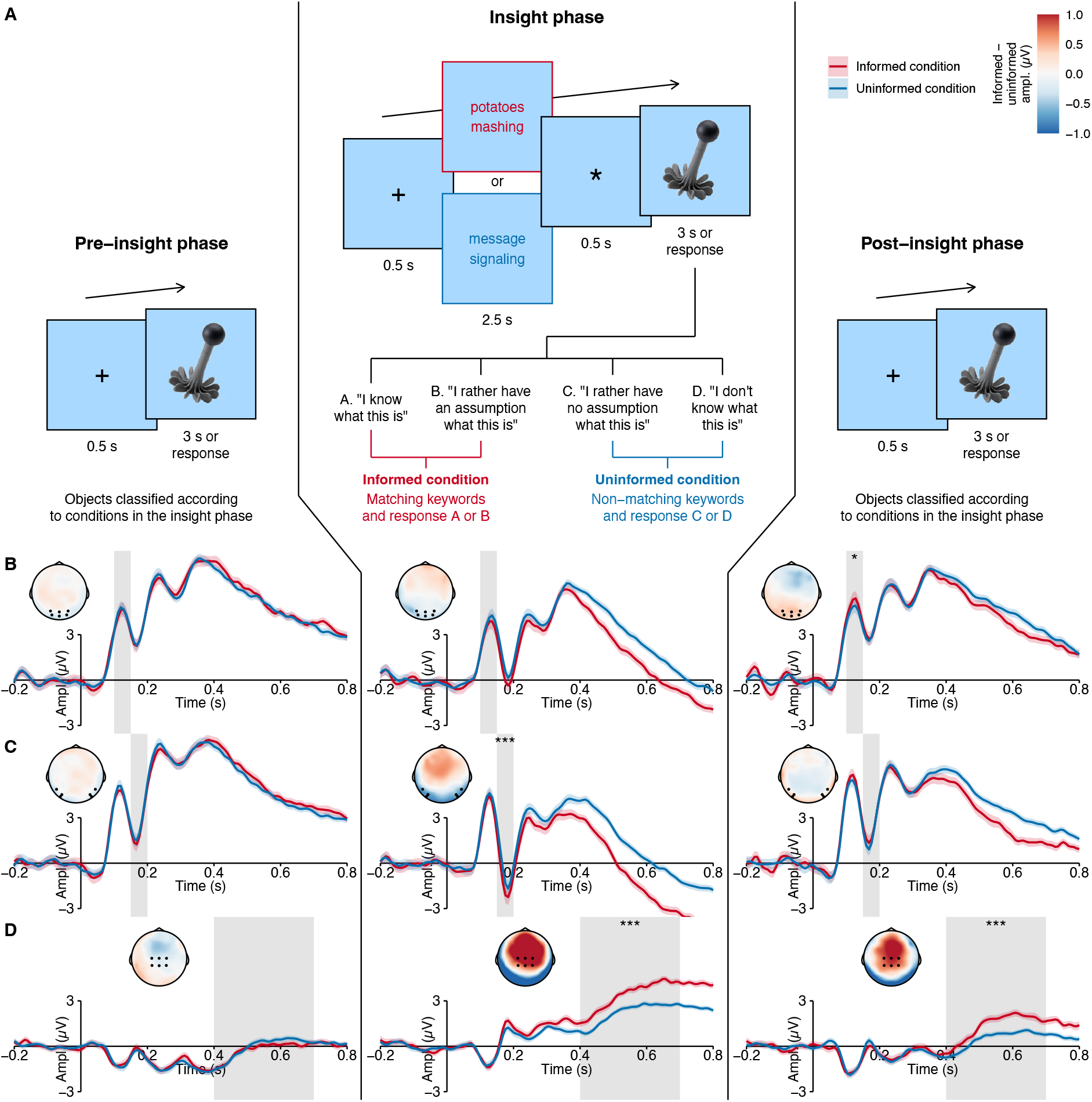
Experimental design and ERP results contrasting semantically informed perception and uninformed perception. ***A***, In the pre-insight phase, participants were presented with 120 unfamiliar objects and indicated if they knew what kind of object they were viewing. In the insight phase, half of these objects were presented with matching keywords (in red color for illustration), leading to semantically informed perception, and the other half with non-matching keywords (in blue color for illustration), leading to uninformed perception. In the post-insight phase, the same objects were presented again without keywords. ***B, C, D***, ERP waveforms and scalp topographies for the P1 component (***B***), for the N170 component (***C***), and for the N400 component (***D***) for objects with semantically informed perception versus uninformed perception within the three different phases. Semantically informed perception was associated with more negative amplitudes in the N170 component during the insight phase, less negative amplitudes in the N400 component during the insight and post-insight phases, and more positive amplitudes in the P1 component during the post-insight phase. Waveform plots show the ERP amplitudes averaged across channels in the regions of interest (P1: PO3, PO4, POz, O1, O2, Oz; N170: P7, P8, PO7, PO8, PO9, PO10; N400: C1, C2, Cz, CP1, CP2, CPz; see black dots in the scalp topographies). Colored ribbons around the ERP waveforms show ± 1 standard error of the mean across participants. Topographies show the difference in ERP amplitudes at all channels on the scalp, averaged across the time windows of interest (P1: 100–150 ms, N170: 150–200 ms, N400: 400–700 ms; see gray areas in the ERP waveforms). Ampl. = amplitude. * *p*_corr_ < .05. *** *p*_corr_ < .001.

**Figure 2.**
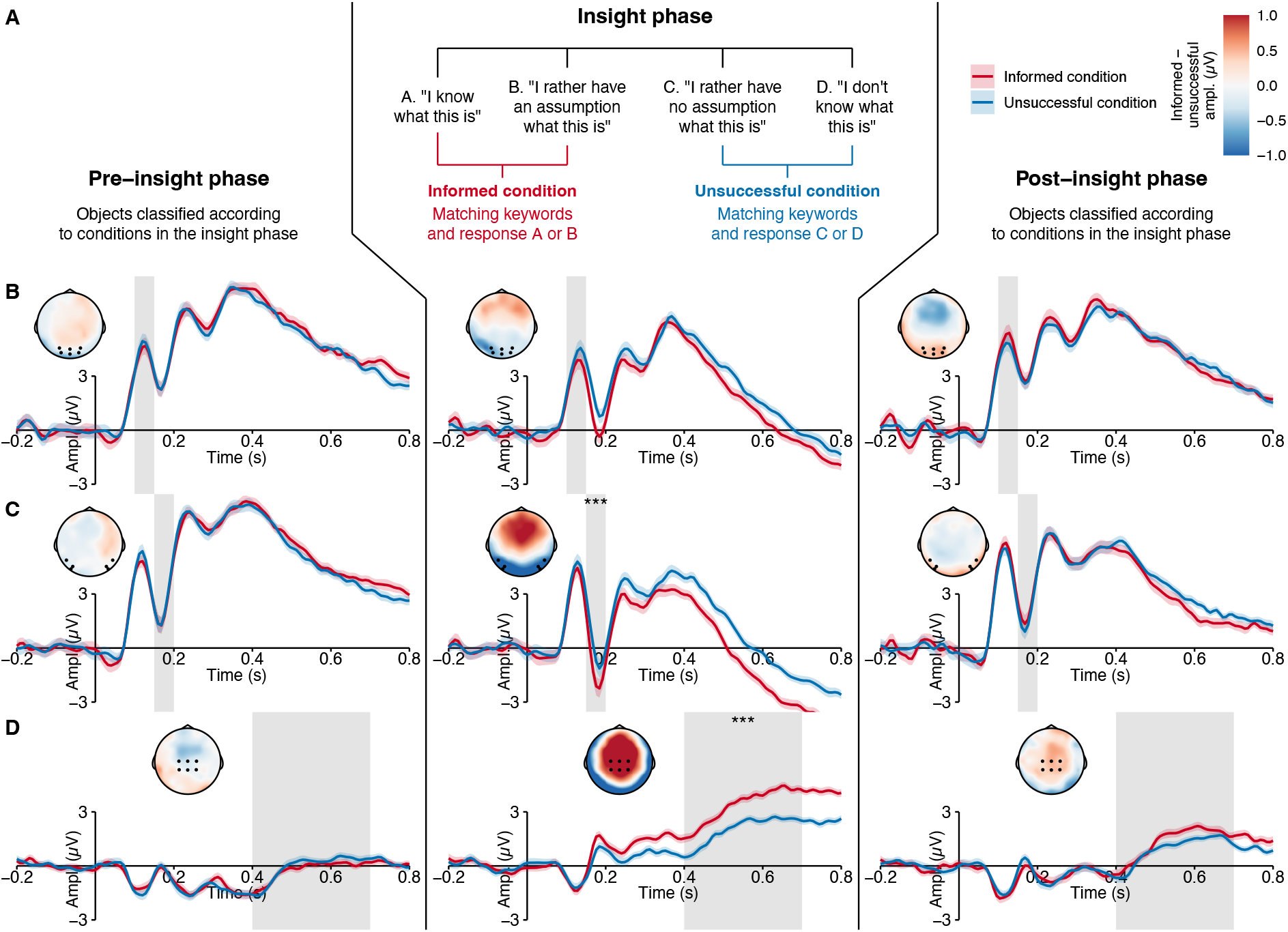
ERP results contrasting semantically informed perception and unsuccessfully informed perception. ***A***, Assignment of objects to conditions based on participants’ response in the insight phase of the experiment to unfamiliar objects presented with matching keywords. ***B, C, D***, ERP waveforms and scalp topographies for the P1 component (***B***), for the N170 component (***C***), and for the N400 component (***D***) for objects with semantically informed perception versus perception within the three different phases. Semantically informed perception was associated with more negative amplitudes in the N170 component and less negative amplitudes in the N400 component during the insight phase. Waveform plots show the ERP amplitudes averaged across channels in the regions of interest (P1: PO3, PO4, POz, O1, O2, Oz; N170: P7, P8, PO7, PO8, PO9, PO10; N400: C1, C2, Cz, CP1, CP2, CPz; see black dots in the scalp topographies). Colored ribbons around the ERP waveforms show ± 1 standard error of the mean across participants. Topographies show the difference in ERP amplitudes at all channels on the scalp, averaged across the time windows of interest (P1: 100–150 ms, N170: 150–200 ms, N400: 400–700 ms; see gray areas in the ERP waveforms). Ampl. = amplitude. *** *p*_corr_ < .001.

### Experimental Design

The main EEG experiment consisted of three phases (see Figure 1A). In the *pre-insight* phase, after written informed consent had been obtained and the EEG had been prepared, all 240 familiar and unfamiliar objects were presented once in random order and without any keywords. Each trial consisted of a fixation cross presented in the middle of the screen for 0.5 s, followed by the presentation of the object until participants made a response or until a time out after 3 s. The inter-trial interval was 0.5 s and participants took a self-timed break after each block of 60 objects. The task, which was kept the same across all three phases of the experiment, was to classify each object using one of four response alternatives: (a) *Ich weiß, was das ist, oder habe eine starke Vermutung* [I know what this is or have a strong assumption], (b) *Ich habe eher eine Vermutung, was das ist* [I rather have an assumption what this is], (c) *Ich habe eher keine Vermutung, was das ist* [I rather have no assumption what this is], or (d) *Ich weiß nicht, was das ist, und habe auch keine Vermutung* [I don’t know what this is and have no assumption]. Participants were asked to respond as quickly and as accurately as possible by pressing one out of four buttons with the index or middle finger of their left or right hand, respectively. The mapping of the rating scale to the four buttons (left to right or right to left) was counterbalanced across participants.

In the *insight* phase, the 120 unfamiliar objects were presented for a second time, now preceded either by matching keywords (leading to semantically informed perception) or by non-matching keywords (leading to uninformed perception). Each trial consisted of a fixation cross presented for 0.5 s, followed by the presentation of the keywords for 2.5 s. Then, an asterisk was presented in the middle of the screen for another 0.5 s, followed by the presentation of the object until a response was made or until a time out after 3 s. The objects were presented in blocks of 30 trials so that within each block there were 15 objects from each of the two experimental conditions and so that objects were heterogeneous in terms of their shape, visual complexity, and functional category (e.g., medical devices, musical instruments).

In the *post-insight* phase, the unfamiliar objects were presented for a third time, with the same trial structure as in the pre-insight phase, that is, without any keywords. The insight and post-insight phases were presented in an interleaved fashion so that after the presentation of one block of 30 objects in the insight phase (with keywords), participants took a self-timed break and continued with the same block of 30 objects in the post-insight phase (without keywords) before moving on to the next block consisting of 30 different objects. They continued like this until all four blocks were completed in both phases. In total, the experiment consisted of 480 trials (120 familiar objects in the pre-insight phase and 120 unfamiliar objects in the pre-insight, insight, and post-insight phases). Participants took approximately 35 minutes to complete the experiment.

### Behavioral Data Analysis

We used participants’ behavioral responses to verify our experimental manipulation and to assign each object to a semantic condition. First, we used participants’ responses from the pre-insight phase (when objects were presented for the first time, without keywords) to make sure that objects were indeed unfamiliar to them. That is, we excluded any objects for which participants responded with “I know what this is” in the pre-insight phase. Next, we used participants’ responses from the insight phase (when objects were presented for the second time, preceded by keywords) to assign the remaining objects to different semantic conditions: Semantically informed perception, uninformed perception, and unsuccessfully informed perception. The semantically informed condition consisted of objects that were presented with matching keywords and for which the participant responded with knowing what the object was or having an assumption. The uninformed condition consisted of objects that were presented with non-matching keywords and for which the participant responded with not knowing what the object was or having rather no assumption. The unsuccessfully informed condition consisted of objects that were presented with matching keywords but for which the participant responded with not knowing what the object was or having rather no assumption. This assignment of objects to semantic conditions (informed, uninformed, or unsuccessfully informed) was carried over from the insight phase to the other two phases (pre-insight and post-insight). This allowed us to test, on the one hand, if the objects differed in important aspects even before any keywords were presented (pre-insight phase) and, on the other hand, if the semantic information acquired in the insight phase had any down-stream effects on the subsequent perception of the objects (post-insight phase).

### EEG Recording and Preprocessing

The continuous EEG was recorded from 62 Ag/AgCl scalp electrodes placed according to the extended 10–20 system (American Electroencephalographic Society, 1991) and referenced online to an external electrode placed on the left mastoid (M1). Two additional external electrodes were placed on the right mastoid (M2) and below the left eye (IO1), respectively. Electrode impedances were kept below 5 kΩ. An online band-pass filter with a high-pass time-constant of 10 s (0.016 Hz) and a low-pass cutoff frequency of 1000 Hz was applied before digitizing the signal at a sampling rate of 500 Hz.

The data were preprocessed offline using custom functions (available at https://github.com/alexenge/hu-neuro-pipeline/tree/v0.6.5) based on the MNE-Python software (Version 1.3.0; Gramfort et al., 2013) in Python (Version 3.8.10; Van Rossum and Drake, 2009). First, the data were downsampled to 125 Hz and re-referenced to the common average of all scalp channels. Next, artifacts resulting from blinks and eye movements were removed via independent component analysis (ICA) on a high-pass filtered copy of the data (cutoff = 1 Hz). A variable number of components per participant were extracted from an initial principal component analysis (PCA) so that they explained at least 99% of the variance in the data (mean = 28.85 components, range 16–40). Then the ICA was fitted based on these components using the FastICA algorithm (Hyvärinen, 1999). Any independent components showing significant correlations with either of two virtual EOG channels (VEOG: IO1 Fp1, HEOG: F9 F10) were removed automatically using MNE-Python’s *find_bads_eog* method. This was the case for an average of 2.38 components per participant (range = 2–4 components).

For the analysis of ERP amplitudes, a zero-phase, non-causal FIR filter with a lower pass-band edge at 0.1 Hz (transition bandwidth: 0.1 Hz) and an upper pass-band edge at 30 Hz (transition bandwidth: 7.5 Hz) was applied. Next, the continuous EEG was segmented into epochs ranging from -500 ms to 1,500 ms relative to the onset of the presentation of each unfamiliar object. These epochs were baseline-corrected by subtracting the average voltage during the interval of -200 ms to 0 ms relative to stimulus onset. Epochs containing artifacts despite ICA, defined as peak-to-peak amplitudes exceeding 150 µV, were removed from further analysis. This led to the exclusion of an average of 11 trials per participant (= 3.1%; range 0–109 trials). Single-trial event-related potentials were computed as the mean amplitude across time windows and regions of interests (ROIs) defined a priori, namely 100–150 ms at channels PO3, PO4, POz, O1, O2, and Oz and for the P1 component, 150–200 ms at channels P7, P8, PO7, PO8, PO9, and PO10 for the N170 component, and 400–700 ms at channels C1, C2, Cz, CP1, CP2, and CPz for the N400 component. We chose a later time window for the N400 component than what is typically used in experiments with verbal materials (e.g., 300–500 ms), in accordance with a previously published data set which showed that the 400–700 ms time window is most robustly associated with the semantic processing of visual objects (Kovalenko et al., 2012).

### Statistical Analysis

The resulting mean ERP amplitudes were analyzed on the single-trial level using linear mixedeffects regression models because these models allow to control for repeated measures of participants and stimuli, while also being robust against an unbalanced number of trials per condition (Bürki et al., 2018; Frömer et al., 2018; Brown, 2021). We computed three models predicting P1, N170, and N400 amplitudes, respectively. All models included the following fixed effects of interest: (a) the phase of the experiment, coded as a repeated contrast (i.e., subtracting the first phase from the second phase and the second phase from the third phase, the intercept being the grand mean across all three phases; Schad et al., 2020), (b) the condition of the object, coded as a custom contrast (i.e., subtracting the uninformed condition from the informed condition and the unsuccessfully informed condition from the informed condition, the intercept being the grand mean across all three conditions), and (c) the twoway interaction of phase and condition. We also added the results from the online pre-rating study (mean cosine distance between rating study-generated keywords and keywords presented in the main experiment) as an additional covariate of no interest (see Online Figure 2). This was to control for the possibility that participants might have been partly familiar with some of the objects and/or able to guess their function from their visual appearance alone. For the random effects, we determined the most parsimonious structure supported by the data using the automatic procedure proposed by Matuschek et al. (2017). This involved starting with a maximal model that contained all random parameters (intercepts, slopes, and correlations) and then iteratively removing terms as long as this did not result in a significant drop in model fit (likelihood ratio test, *p* value cutoff = .20; see Online Results 1 for the final model syntax and model outputs). All models were fitted in R (Version 4.2.1; R Core Team, 2022) using the lme4 package (Version 1.1.30; Bates et al., 2015) with the optimizer function *bobyqa* and a maximum of 10^6^ iterations for maximum likelihood estimation. The model selection algorithm via likelihood ratio tests was performed using the buildmer package (Version 2.7; Voeten, 2022).

To investigate if semantically informed perception had an influence on the ERPs within each phase of the experiment, we calculated pairwise comparisons contrasting the semantically informed condition against the uninformed condition within the pre-insight, insight, and post-insight phases. In the same way, we computed pairwise comparisons contrasting the semantically informed condition against the unsuccessfully informed condition. This was done using the emmeans package (Version 1.8.2; Lenth, 2022) and with Bonferroni correction for the three phases of the experiment. All *p* values were computed by approximating the relevant denominator degrees of freedom using Satterthwaite’s method as implemented in the lmerTest package (Version 3.1.3; Kuznetsova et al., 2017).

### Time-Frequency Analysis

For our exploratory analysis of event-related power, we first created new epochs from the ICA-corrected but unfiltered raw data. Epochs that were marked as bad for the ERP analysis (i.e., with peak-to-peak amplitudes exceeding 150 µV) were also removed from the time-frequency analysis. The remaining epochs were convolved with a family of Morlet wavelets, increasing linearly in their frequency from 4 Hz to 40 Hz in steps of 1 Hz and in their width from 2 cycles to 20 cycles in steps of 0.5 cycles. To adjust for the typical 1/*f* shape of the EEG, the power values were transformed into percent signal change by first subtracting and then dividing by the average power at each frequency over the entire epoch (Grandchamp and Delorme, 2011). We then performed baseline correction by subtracting the average power during the pre-stimulus interval from -450 ms to -50 ms s relative to object onset.

For statistical analysis of event-related power, we conducted cluster-based permutation tests (Maris and Oostenveld, 2007), separately for each of the three phases of the experiment (pre-insight, insight, and post-insight). First, we averaged trials belonging to the same condition and then subtracted these average responses from one another to compute the difference between the semantically informed condition and each of two control conditions (uninformed or unsuccessfully informed) for each participant. We then conducted one-sample *t* tests in a mass-univariate fashion and grouped significant results (cluster-forming threshold *p* < .05) into clusters if they occurred at neighboring time points, frequencies, or channels. Neighboring channels were defined using the Delaunay triangulation based on 2D electrode locations as implemented in MNE-Python. To obtain family-wise error-corrected *p* values at the cluster level, we compared the cluster mass (i.e., the sum of the *t* values) of each observed cluster to an empirical null distribution of cluster masses obtained from 5,000 permutations with random sign flips. Clusters were considered to be statistically significant if their mass exceeded the 98.33^th^ percentile of this distribution (i.e., cluster-level threshold *p* < .05, Bonferroni-corrected for three phases of the experiment).

### Data and Code Accessibility

The EEG data are available upon reasonable request from the corresponding author because we had not asked participants for their consent to make the data publicly available. The experimental stimuli (object images and keywords) and the code for data analysis are available at https://doi.org/10.17605/osf.io/uksbc.

In addition to the software mentioned above, our code relies on the tidyverse set of R packages (Version 1.3.2; Wickham et al., 2019) for data wrangling, the ggplot2 (Version 3.3.6; Wickham, 2016), cowplot (Version 1.1.1; Wilke, 2020), and eegUtils (Version 0.7.0; Craddock, 2022) packages for visualization, the papaja package (Version 0.1.1; Aust and Barth, 2022) for statistical reporting, and the LSAfun package (Version 0.6.3; Günther et al., 2015) for the latent semantic analysis of the online rating study data. We used the workflow developed by Peikert and Brandmaier (2021) to ensure the long-term reproducibility of our analysis pipeline.

## Results

### Behavioral Data

Based on the experimental manipulation (matching or non-matching keywords) and each individual participant’s behavioral response (positive or negative regarding knowing what the object was), we assigned 26.67 ± 10.51 (mean ± standard deviation) objects to the semantically informed condition (i.e., matching keywords and positive response), 48.98 ± 7.71 objects to the uninformed condition (i.e., non-matching keywords and negative response), and 26.77 ± 10.48 objects to the unsuccessfully informed condition (i.e., matching keywords but negative response). This assignment of objects to conditions was based solely on the insight phase of the experiment, when objects were presented with keywords for the first time, and carried over to analyze the data from the pre-insight and post-insight phases as well. The remaining objects were excluded due to an implausible response pattern (i.e., non-matching keywords but positive response; 5.29 ± 5.78 objects), due to being known to the participant in the pre-insight phase (i.e., before any keywords had been presented; 7.10 ± 5.00 objects), or because of technical errors (i.e., reaction times of 0 ms recorded by the presentation software, 5.19 ± 5.84 objects).

### Event-Related Potentials

Averaged across conditions, P1, N170, and N400 amplitudes differed as a function of the phase of the experiment (pre-insight, insight, or post-insight), all *f* s > 14.28, all *p*s < .001. In addition, N400 amplitudes differed as a function of the condition (semantically informed, uninformed, or unsuccessfully informed), averaged across the three phases of the experiment, *f* (2, 14,180) = 22.91, *p* < .001. Crucially, the phase × condition interaction was significant for all three ERP components, all *f* s > 2.54, all *p*s < .038. To answer our main research question, we decomposed these interactions into pairwise comparisons between semantic conditions within each of the three phases of the experiment.

In the pre-insight phase, when objects were unfamiliar to participants and presented without keywords, no differences emerged between the semantically informed condition and the uninformed condition in any of the the ERP components, all |*t*|s < 0.86, all *p*s > .999 (see Figure 1B–D). Likewise, there were no differences between the semantically informed condition and the unsuccessfully informed condition in any of the the ERP components, all |*t*|s < 0.55, all *p*s > .999 (see Figure 2B–D). This was expected given that the keywords that would provide additional semantic information had not yet been presented, and given that the assignment of objects to conditions was counterbalanced across participants as to control for low-level visual differences.

In the insight phase, half of the unfamiliar objects were presented with matching keywords, leading to semantically informed perception, and the other half were presented with non-matching keywords, keeping the perception of the objects semantically uninformed. Semantically informed perception in this phase was associated with enlarged (i.e., more negative) amplitudes in the N170 component, *b* = -0.64 µV, *t*(14,194) = -3.63, *p*_corr_ = .001, and reduced (i.e., less negative) amplitudes in the N400 component, *b* = 1.09 µV, *t*(14,156) = 7.17, *p*_corr_ < .001. There was no reliable difference between conditions in the P1 component, *b* = -0.27 µV, *t*(14,138) = -1.49, *p*_corr_ = .406. The same effects were found when comparing objects with semantically informed perception to those objects that were also presented with matching keywords but for which participants still indicated not knowing what the object was (i.e., the unsuccessfully informed condition). Compared to these objects, semantically informed perception was associated with enlarged (i.e., more negative) amplitudes in the N170 component, *b* = -0.90 µV, *t*(14,203) = -4.37, *p*_corr_ = < .001, and reduced (i.e., less negative) amplitudes in the N400 component, *b* = 1.22 µV, *t*(13,925) = 7.00, *p*_corr_ < .001, while there was no reliable difference in the P1 component, *b* = -0.39 µV, *t*(13,860) = -1.84, *p*_corr_ = .196.

In the post-insight phase, the unfamiliar objects were presented for a third time and without any keywords, mirroring the pre-insight phase. As in the insight phase, semantically informed perception was associated with reduced (i.e., less negative) amplitudes in the N400 component, *b* = 0.80 µV, *t*(14,031) = 5.23, *p*_corr_ < .001, while the effect in the N170 component did not recur, *b* = 0.26 µV, *t*(13,742) = 1.46, *p*_corr_ = .431. Instead, the P1 component was significantly enlarged (i.e., more positive) in response to objects for which semantically informed perception had taken place, *b* = 0.46 µV, *t*(13,047) = 2.52, *p*_corr_ = .036. These effects did not replicate when comparing the semantically informed condition to the unsuccessfully informed condition, with no reliable differences in the P1, N170, or N400 components, all |*t*|s < 2.17, all *p*s > .090.

### Event-Related Power

In an exploratory time-frequency analysis, we checked for differences in event-related power between semantically informed perception and uninformed perception within each of the three phases of the experiment. Cluster-based permutation tests (Maris and Oostenveld, 2007) revealed no significant clusters in the pre-insight phase (see Online Figure 3), all *p*s > .789, but one significant cluster in the insight phase (see Figure 3), *p*_cluster_ = .002, and one significant cluster in the post-insight phase (see Figure 4), *p*_cluster_ = .002. These two significant clusters in the insight and post-insight phases were similar in their direction, latency, frequency range, and topographic distribution. Both clusters had a negative sign, started around 600 ms after object onset (but see Sassenhagen and Draschkow, 2019) and continued all the way until the end of the analyzed period at 1,400 ms. They spanned a broad range of frequencies in the alpha and lower beta range as well as a broad set of channels, but appeared to be most focal at around 15 Hz and parietal channels. Thus, semantically informed perception seems to alter not only early, evoked activity (see Event-Related Potentials above) but also later, induced activity, in the form of a reduction of post-stimulus power at parietal channels in the range of alpha and lower beta frequencies.

**Figure 3.**
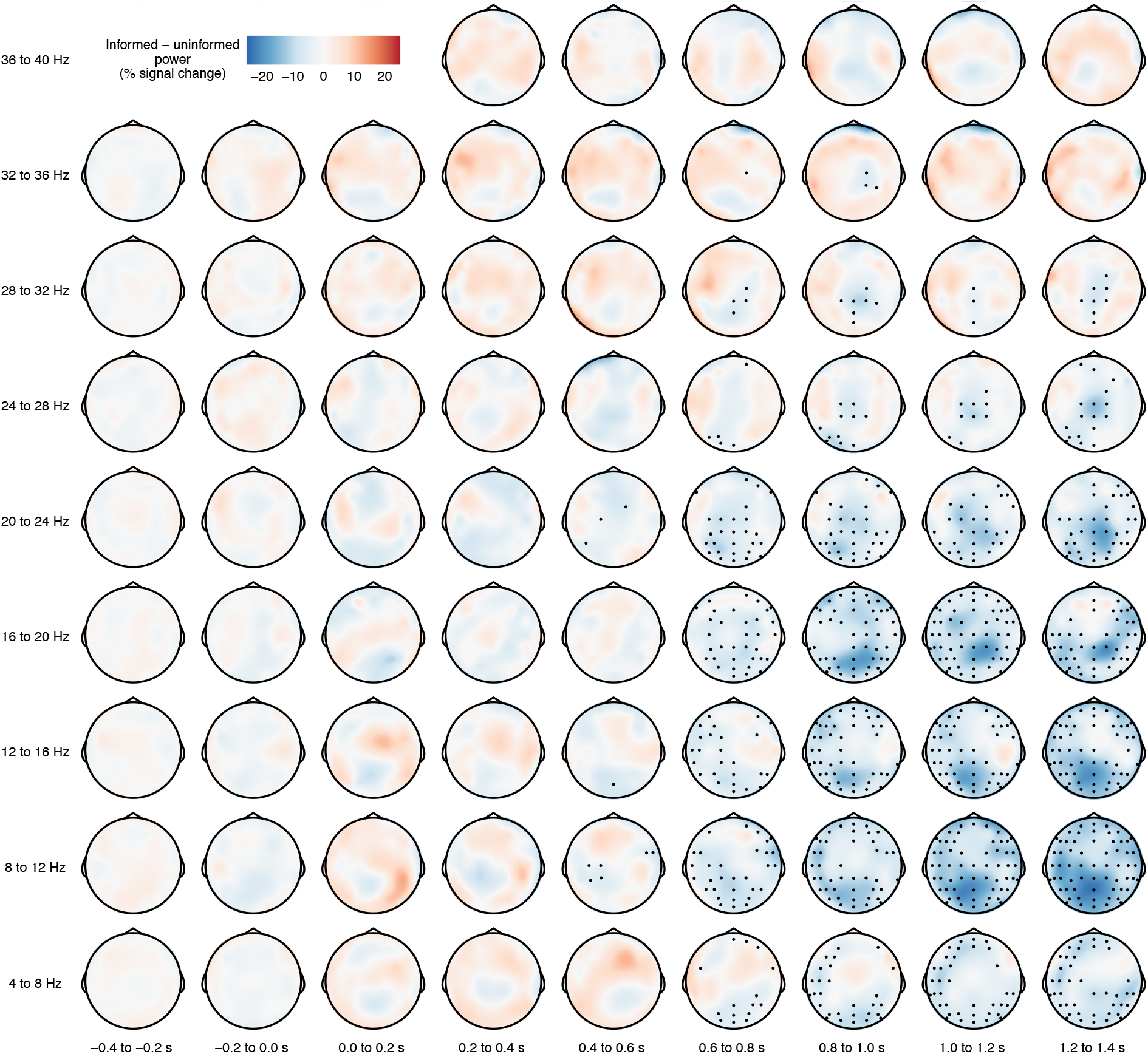
Time-frequency results for the insight phase. Each topographic plot shows the difference in event-related power (in units of percent signal change) between the semantically informed condition and the uninformed condition, grand-averaged across participants. Black dots highlight EEG channels that were part of a cluster for which this difference was statistically significant (*p*_cluster_ = .002).

**Figure 4.**
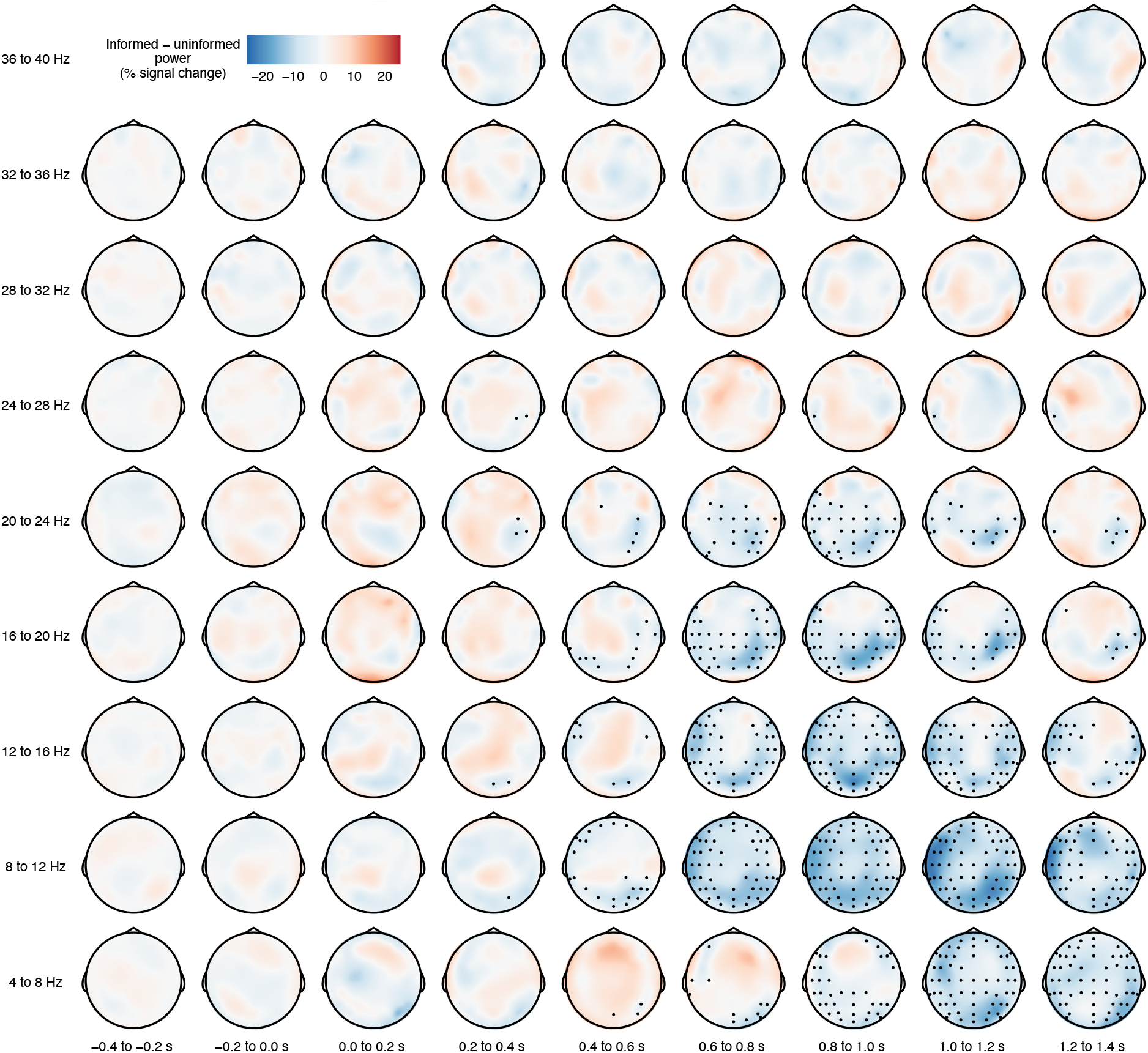
Time-frequency results for the post-insight phase. Each topographic plot shows the difference in event-related power (in units of percent signal change) between the semantically informed condition and the uninformed condition, grand-averaged across participants. Black dots highlight EEG channels that were part of a cluster for which this difference was statistically significant (*p*_cluster_ = .002).

We repeated this analysis to look for differences between semantically informed perception and unsuccessfully informed perception within each of the three phases of the experiment. There were no significant clusters in the pre- or post-insight phases (see Online Figures 4 and 6), all *p*s > .216 but one significant cluster in the insight phase (see Online Figure 5), *p*_cluster_ = .016. This cluster was similar to the ones described above in its latency, frequency range, and topographic distribution.

## Discussion

We found that providing participants once with semantic information about previously unfamiliar objects instantly led to enlarged (i.e., more negative) ERP amplitudes in the N170 component and reduced (i.e., less negative) ERP amplitudes in the N400 component. When the same objects were presented again, the N400 component remained reduced and the P1 component was now enlarged (i.e., more positive) in response to objects that had previously triggered semantically informed perception. An exploratory time-frequency analysis revealed that semantically informed perception was accompanied by a late reduction in event-related power in the alpha and lower beta ranges.

The N400 effect indicates that acquiring an understanding of the object lessened participants’ demand for semantic processing (Kutas and Federmeier, 2011). It replicates previous work showing larger N400 amplitudes for pictures when they are difficult to understand in and of themselves (e.g., Supp et al., 2005; Abdel Rahman and Sommer, 2008) or difficult to integrate into the preceding context (e.g., Barrett and Rugg, 1990; Ganis et al., 1996; Hirschfeld et al., 2011). The latency of this effect suggests a post-perceptual locus in the semantic system.

Our exploratory time-frequency analysis revealed a late (> 600 ms) reduction in power in the alpha and lower beta ranges (approx. 8–20 Hz). Like the N400 effect, this occurred as soon as participants had received the semantic information and recurred once the objects were re-encountered without any semantic information. Reductions in alpha/beta power have been shown to correlate with clearer representations of stimulus specific information, as measured using representational similarity analysis (Griffiths et al., 2019), and with successfully forming new semantic memories (Hanslmayr et al., 2009). This may be due to a dampening of alpha/beta oscillations which creates favorable conditions for high-level cortical information processing and encoding.

In contrast to the N400 and event-related power, the N170 was modulated only on those initial trials on which the relevant semantic information was presented directly before the object. It therefore constitutes an online marker of semantic insight, that is, of participants suddenly understanding the visual objects in the light of the information provided by the keywords. The N170 is associated with the holistic perception of faces (Sagiv and Bentin, 2001; Eimer et al., 2011) and other stimuli of visual expertise (Tanaka and Curran, 2001; Rossion et al., 2002), that is, it is sensitive to factors that go beyond structural encoding and categorical perception (Thierry et al., 2007a, 2007b; Dering et al., 2011). It being enlarged may reflect that the semantic information made participants experience the configuration of the visual features of the objects in a new and meaningful way. This is supported by previous findings of enlarged N170 amplitudes for scrambled face stimuli after participants had been shown the original version of the face (Bentin and Golland, 2002), as well as for line drawings of meaningful objects as compared to non-objects (Beaucousin et al., 2011). Together, this suggests an online impact of meaningfulness on the higher-level perception of visual objects, integrating across their visual features.

The P1, unlike the N400 and N170, was modulated by semantic information only one trial after this information had been obtained. This replicates previous studies showing modulations of the P1 when participants had learned meaningful information about previously unfamiliar objects (Abdel Rahman and Sommer, 2008; Maier and Abdel Rahman, 2018, 2019; Samaha et al., 2018; Weller et al., 2019). The present study adds that the P1 effect does not take an extensive learning history to develop; instead, it can be observed as soon as one trial after semantic insight has happened. Because the P1 is associated with lower-level sensory processing (Johannes et al., 1995; Pratt, 2011; Luck, 2014), we take its susceptibility to semantic information as an indicator that knowledge about the function of an object can change how we perceive its low-level features (Athanasopoulos and Casaponsa, 2020).

The P1 and N170 were modulated in different phases of our study, suggesting that they reflect different aspects of top-down processing with different time courses and neuroanatomical implementations. The time course of the N170 is consistent with a top-down influence of non-visual areas in the prefrontal and temporo-parietal cortices on visual areas, whereas modulations of the P1 component may reflect recurrent processing within the visual system (Wyatte et al., 2014). Here we could show that the former pathway seems to be able to convey semantic information instantaneously (i.e., within the same trial), whereas the latter pathway seems to take at least one additional encounter with the object to emerge. While the limited spatial resolution of the EEG precludes localization, there is converging fMRI and psychophysical evidence that semantic information can feed back into areas in the lateral occipital cortex (LOC) as well as early retinotopic cortex (areas V1, V2, and V3; Hsieh et al., 2010; Clarke et al., 2016; Teufel et al., 2018), consistent with the neural generators of the N170 and P1 in the ERP.

The top-down modulation of visual ERPs by semantic information challenges a modular view of visual perception (Fodor, 1983; Pylyshyn, 1999). Proponents of this view have pointed out important shortcomings of previous studies that had claimed to demonstrate top-down effects of cognition on perception (Machery, 2015; Firestone and Scholl, 2016). We addressed as many of these shortcomings as possible: We ensured that there were no visual differences between conditions (with counterbalancing and the pre-insight phase as a negative control), we used ERPs as an objective and time-resolved measure to disentangle perceptual and post-perceptual effects, and we reduced response and demand biases by keeping the manipulation (i.e., matching or non-matching keywords) obscure to participants and by including well-known objects as filler stimuli.

One could argue that the effects presented here might be reducible to more basic mechanisms such as semantic priming. Indeed, the keywords that were presented before each object in the semantically informed condition were chosen such that they matched the function of the object and often had a direct relationship to certain visual features of the object. This might have induced semantic priming, which is supported by the reduction in N400 amplitudes (e.g., Bentin et al., 1985; Kellenbach et al., 2000). However, there are at least three arguments why semantic priming cannot account for our main findings, that is, the influence of semantic information on the P1 and N170. First, for both components, ERP amplitudes were enlarged (i.e., more positive for the P1 and more negative for the N170) during semantically informed perception, whereas semantic priming typically leads to reduced ERP amplitudes. Second, these effects were not just observed when comparing semantically informed perception (with matching keywords) and uninformed perception (with non-matching keywords), but also when comparing semantically informed perception with unsuccessfully informed perception. In the latter case, all objects were preceded by keywords that matched the function of the object to a similar degree, making semantic priming less likely. Third, the results from an online rating study (see Materials and Methods; Online Figure 1) indicated that people by and large did not spontaneously associate the unfamiliar objects with a particular function, and also allowed us to statistically control for the closeness of peoples’ guesses and the true function of each object.

A theoretical framework that would explicitly predict the observed P1 and N170 effects in our study is lacking at present. However, the effects are consistent with the reverse hierarchy theory (Ahissar and Hochstein, 2004), which posits that objects first enter visual consciousness at an abstract, conceptual level. Once this initial “vision at a glance” has taken place, feedback connections to earlier layers of the visual system are being accessed to extract the relevant lower-level features (“vision with scrutiny”). This reverse trajectory down the visual hierarchy may explain (a) the semantically induced changes to the fMRI signal in LOC and retinotopic cortex (e.g., Hsieh et al., 2010) as well as (b) the modulations of early visual ERP components observed in the present study and others (e.g., Abdel Rahman and Sommer, 2008; Maier et al., 2014; Samaha et al., 2018). An important role of top-down mechanisms for object recognition is also posited by theories of predictive coding and Bayesian inference (e.g., Yuille and Kersten, 2006; Xu and Tenenbaum, 2007; Clark, 2013; Panichello et al., 2013; Lupyan, 2015). Despite the theoretical advances, detailed descriptions of these top-down effects at the algorithmic and implementational levels remain a challenge for future work.

Taken together, the present study provides preliminary evidence that whenever we receive semantic information about a previously unfamiliar object, this information has an immediate influence on our visual processing of this object. The immediacy of this influence is remarkable in at least two different ways: First, it does not require an extensive learning history but can instead be observed within the same trial in which the information has been presented and/or a single trial later. Second, the time course of this influence suggests that it manifests itself not only at later, post-perceptual stages (> 400 ms), typically associated with semantic processing, but also at much earlier stages within the first 200 ms, associated with visual perception itself.

