## Supplementary material for "Instant Effects of Semantic Information on Visual Perception": Online supplementary information

Alexander Enge, Franziska Süß, & Rasha Abdel Rahman

**Online Table 1.** ***Unfamiliar Object Stimuli***

| Stimulus | ID | Matching keywords | Non-matching keywords |
|---|---|---|---|
| 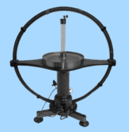 | 1 | elektrische Spannung, prüfen<br>[electric current, measuring] | Makkaroni, formen<br>[macaroni, forming] |
| 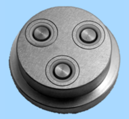 | 2 | Makkaroni, formen<br>[macaroni, forming] | Kuh, vom Zaun abhalten<br>[cow, keeping away from the fence] |
| 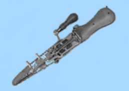 | 3 | Knochen, sägen<br>[bones, sawing] | Streckenmaß, Sonnenlicht nutzen<br>[distance measure, using sunrays] |
| 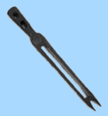 | 4 | Unkraut, jäten<br>[weed, removing] | Uhr, mit Wärme betreiben<br>[clock, operating with heat] |
| 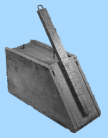 | 5 | Mausefalle, zuschnappen<br>[mousetrap, snap-shutting] | Brillenglas, zuschneiden<br>[eyeglass lens, cutting] |
| 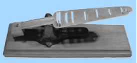 | 6 | Goldmünzen, wiegen<br>[gold coin, weighing] | Farbe, vom Fenster abschleifen<br>[paint, scraping off the window] |
| 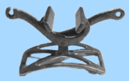 | 7 | Knie, fixieren<br>[knee, fixating] | Buchstaben, tippen<br>[letter, typing] |
| 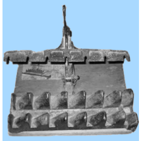 | 8 | Eierkarton, pressen<br>[egg carton, pressing] | Tabak, zermahlen<br>[tobacco, pulverize] |
| 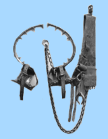 | 9 | Baum, erklettern<br>[tree, climbing] | Rotation, Ladung erzeugen<br>[rotation, creating electric charge] |
| 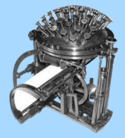 | 10 | Buchstaben, tippen<br>[letter, typing] | Pferdehuf, Halt geben<br>[horse hoof, giving grip] |
| 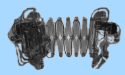 | 11 | Akkordeon, spielen<br>[accordion, playing] | Seil, schneiden<br>[rope, cutting] |

Table 1 continued

| Stimulus | ID | Matching keywords | Non-matching keywords |
|---|---|---|---|
| 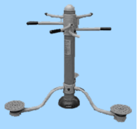 | 12 | Körper, trainieren [body, training] | Buch, offen halten [book, binding] |
| 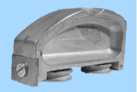 | 13 | Farbe, vom Fenster abschleifen [paint, scraping off the window] | von Hand, zentrifugieren [by hand, centrifugating] |
| 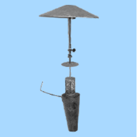 | 14 | Außenbereich, heizen [outdoor area, heating] | Rasierklinge, schärfen [razor blade, sharpening] |
| 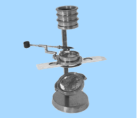 | 15 | Pflanzenteile, vergrößern [plant parts, magnifying] | Katzenklo, sich selbst reinigen [litter box, self-cleaning] |
| 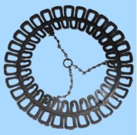 | 16 | Krawatten, aufhängen [necktie, hanging up] | Zeichnungen, vermessen [drawings, measuring] |
| 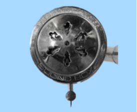 | 17 | Schallplatte, abtasten [vinyl record, reading] | Ball, katapultieren [ball, catapulting] |
| 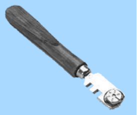 | 18 | Glas, schneiden [glass, cutting] | Eier, wiegen [eggs, weighing] |
| 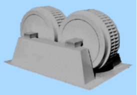 | 19 | Briketts, pressen [briquette, pressing] | Narkosemittel, abgeben [anesthetic, administering] |
| 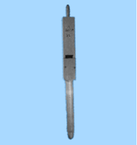 | 20 | Orgelton, erzeugen [organ sound, making] | Bandage, rollen [bandage, rolling] |
| 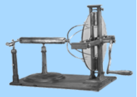 | 21 | Spannung, erzeugen [electric current, making] | Fußstütze, reiten [footrest, horseback riding] |
| 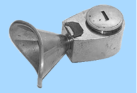 | 22 | Narkosemittel, abgeben [anesthetic, administering] | Nüsse, aufbrechen [nuts, cracking] |
| 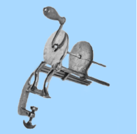 | 23 | Bandage, rollen [bandage, rolling] | Schnee, rodeln [snow, sleighing] |
| 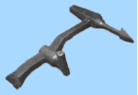 | 24 | Weinfass, Loch einschlagen [wine barrel, smashing a hole] | Tier, einfangen [animal, trapping] |

Table 1 continued

| Stimulus | ID | Matching keywords | Non-matching keywords |
|---|---|---|---|
| 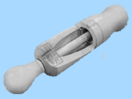 | 25 | Flaschenkorken, einführen<br>[bottle cork, inserting] | Pferd, im Moor laufen<br>[horse, walking in the moor] |
| 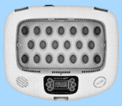 | 26 | Automat, Eier ausbrüten<br>[machine, incubating eggs] | Windgeschwindigkeit, messen<br>[wind speed, measuring] |
| 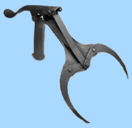 | 27 | Korngarbe, greifen<br>[sheaf, grabbing] | Treibhaus, heizen<br>[glass house, heating] |
| 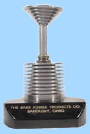 | 28 | Brief, wiegen<br>[letter, weighing] | Korngarbe, greifen<br>[sheaf, grabbing] |
| 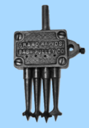 | 29 | Löcher, bohren<br>[hole, drilling] | Sonnenlicht, Intensität messen<br>[sunlight, measure intensity] |
| 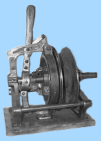 | 30 | Angelschnur, kurbeln<br>[fishing line, winding] | Glaskörper, musizieren<br>[glass body, making music] |
| 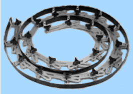 | 31 | Kurven, malen<br>[curve, drawing] | Nussöl, pressen<br>[nut oil, pressing] |
| 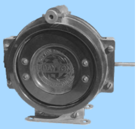 | 32 | Radiofrequenz, einstellen<br>[radio frequency, tuning] | Erektion, helfen<br>[erection, helping] |
| 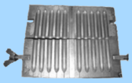 | 33 | Zäpfchen, pressen<br>[suppository, pressing] | Unkraut, jäten<br>[weed, removing] |
| 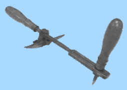 | 34 | Fass, öffnen<br>[barrel, opening] | Pflanzenteile, vergrößern<br>[plant parts, magnifying] |
| 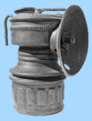 | 35 | Bergbaustollen, beleuchten<br>[mining tunnel, lightening] | Knie, fixieren<br>[knee, fixating] |
| 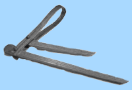 | 36 | Kuh, vom Zaun abhalten<br>[cow, keeping away from the fence] | Radiofrequenz, einstellen<br>[radio frequency, tuning] |
| 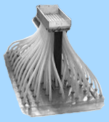 | 37 | Saatgut, gleichmäßig aussäen<br>[seeds, sowing evenly] | Fass, öffnen<br>[barrel, opening] |

Table 1 continued

| Stimulus | ID | Matching keywords | Non-matching keywords |
|---|---|---|---|
| 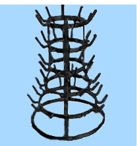 | 38 | Flaschen, trocknen<br>[bottles, drying] | Bleistift, anspitzen<br>[pencil, sharpening] |
| 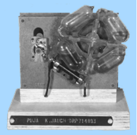 | 39 | Uhr, mit Wärme betreiben<br>[clock, operating with heat] | Körper, untersuchen<br>[body, training] |
| 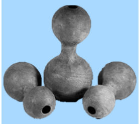 | 40 | Tonpott, trommeln<br>[clay pot, drumming] | Piano, stimmen<br>[piano, tuning] |
| 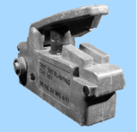 | 41 | Sprengstoffexplosion, auslösen<br>[dynamite explosion, triggering] | Fisch, wiegen<br>[fish, weighing] |
| 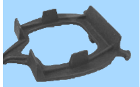 | 42 | Pferdehuf, Halt geben<br>[horse hoof, giving grip] | Zäpfchen, pressen<br>[suppository, pressing] |
| 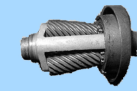 | 43 | Bleistift, anspitzen<br>[pencil, sharpening] | Angel, Köder markieren<br>[fishing rod, marking bait] |
| 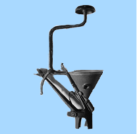 | 44 | Nussöl, pressen<br>[nut oil, pressing] | Zeichen, einbrennen<br>[marks, burning in] |
| 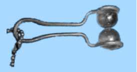 | 45 | Rasierklinge, schärfen<br>[razor blade, sharpening] | Kurven, malen<br>[curve, drawing] |
| 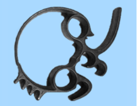 | 46 | heiße Platten, anheben<br>[hot plates, lifting] | Kleidung, im Eimer waschen<br>[clothes, washing in a bucket] |
| 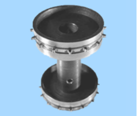 | 47 | Film, aufspulen<br>[film roll, winding] | Mund, offen halten<br>[mouth, keeping open] |
| 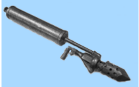 | 48 | Waffe, entflammen<br>[weapon, inflaming] | Uhrzeit, anzeigen<br>[time, displaying] |
| 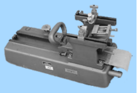 | 49 | Mikroskop-Proben, schneiden<br>[microscopic samples, slicing] | heiße Platten, anheben<br>[hot plates, lifting] |
| 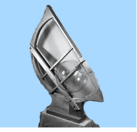 | 50 | Glaskörper, musizieren<br>[glass body, making music] | Kork, flach pressen<br>[cork, pressing flat] |

Table 1 continued

| Stimulus | ID | Matching keywords | Non-matching keywords |
|---|---|---|---|
| 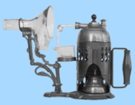 | 51 | Dampf, zerstäuben<br>[steam, spraying] | Buch, binden<br>[book, opening] |
| 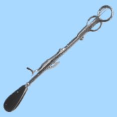 | 52 | Mandeln, operieren<br>[kidneys, operating] | Film, aufspulen<br>[film roll, winding] |
| 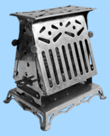 | 53 | Toastbrot, rösten<br>[toast, roasting] | Autobatterie, Spannung testen<br>[car battery, measuring voltage] |
| 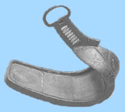 | 54 | Fußstütze, reiten<br>[footrest, horseback riding] | Stromstärke, messen<br>[current, measuring] |
| 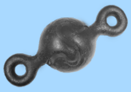 | 55 | Türgelenk, Feuer überstehen<br>[door hinge, surviving fires] | Botschaft, telegrafieren<br>[message, telegraphing] |
| 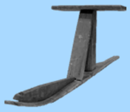 | 56 | Schnee, rodeln<br>[snow, sleighing] | Dampf, zerstäuben<br>[steam, spraying] |
| 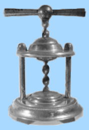 | 57 | Nüsse, aufbrechen<br>[nuts, cracking] | Ziegelsteine, formen<br>[bricks, forming] |
| 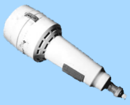 | 58 | Radiergummi, mit Strom betreiben<br>[eraser, operating with electricity] | Münzen, aufbewahren<br>[coins, storing] |
| 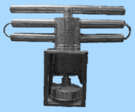 | 59 | Treibhaus, heizen<br>[glass house, heating] | Messer, schleifen<br>[knife, sharpening] |
| 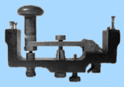 | 60 | Botschaft, telegrafieren<br>[message, telegraphing] | Toastbrot, rösten<br>[toast, roasting] |
| 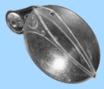 | 61 | Angel, Köder markieren<br>[fishing rod, marking bait] | Baum, erklettern<br>[tree, climbing] |
| 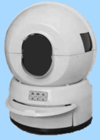 | 62 | Katzenklo, sich selbst reinigen<br>[litter box, self-cleaning] | Tankfüllstand, messen<br>[fuel tank level, gauging] |
| 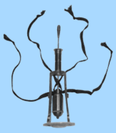 | 63 | Körper, untersuchen<br>[body, examining] | Schuhe, auf Eis laufen<br>[shoes, walking on ice] |

Table 1 continued

| Stimulus | ID | Matching keywords | Non-matching keywords |
|---|---|---|---|
| 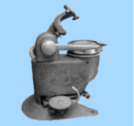 | 64 | Brillenglas, zuschneiden [eyeglass lens, cutting] | Saatgut, gleichmäßig aussäen [seeds, sowing evenly] |
| 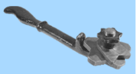 | 65 | Draht, wickeln [wire, winding] | Bergbaustollen, beleuchten [mining tunnel, lightening] |
| 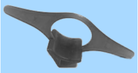 | 66 | Buch, offen halten [book, opening] | Sprengstoffexplosion, auslösen [dynamite explosion, triggering] |
| 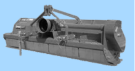 | 67 | Feldanbau, häckseln [harvest, chopping] | Draht, wickeln [wire, winding] |
| 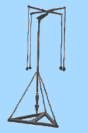 | 68 | Schnurlot, absenken [plummet, letting down] | Korken, formen [bottle cork, forming] |
| 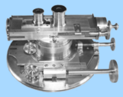 | 69 | Sternenbilder, vermessen [stellar constellation, measuring] | Schnurlot, absenken [plummet, letting down] |
| 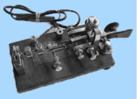 | 70 | Nachricht, morsen [message, signaling] | Feldanbau, häckseln [harvest, chopping] |
| 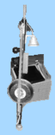 | 71 | Musikgerät, stampfen [musical instrument, stamping] | Goldmünzen, wiegen [gold coin, weighing] |
| 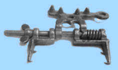 | 72 | Schuhe, auf Eis laufen [shoes, walking on ice] | Geschwindigkeit, ermitteln [speed, determining] |
| 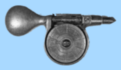 | 73 | Geschwindigkeit, ermitteln [speed, determining] | Mausefalle, zuschnappen [mousetrap, snap-shutting] |
| 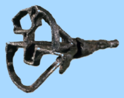 | 74 | Zeichen, einbrennen [marks, burning in] | Luftdruck, messen [air pressure, measuring] |
| 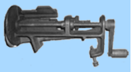 | 75 | Tabak, zermahlen [tobacco, pulverize] | Flaschen, trocknen [bottles, drying] |
| 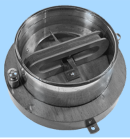 | 76 | Stromstärke, messen [current, measuring] | Mandeln, operieren [kidneys, operating] |

Table 1 continued

| Stimulus | ID | Matching keywords | Non-matching keywords |
|---|---|---|---|
| 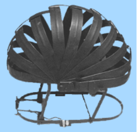 | 77 | Tier, einfangen [animal, trapping] | Maiskolben, entkörnen [corncob, removing grains] |
| 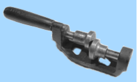 | 78 | Messer, schleifen [knife, sharpening] | Schritte, vergrößern [steps, extending] |
| 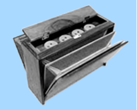 | 79 | Leierkasten, klingen [barrel organ, making sounds] | Fass, anheben [barrel, lifting] |
| 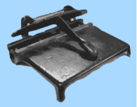 | 80 | Buch, binden [book, binding] | Löcher, bohren [hole, drilling] |
| 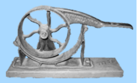 | 81 | Kork, flach pressen [cork, pressing flat] | Elektroschock, spielen [electric shock, playing] |
| 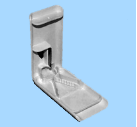 | 82 | Tabletten, zerteilen [pill, splitting] | Blumentopf, sich selbst wässern [flowerpot, self-watering] |
| 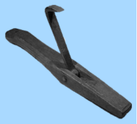 | 83 | Fass, anheben [barrel, lifting] | Mikroskop-Proben, schneiden [microscopic samples, slicing] |
| 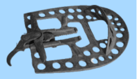 | 84 | Pferd, im Moor laufen [horse, walking in the moor] | Luft, abpumpen [air, pumping out] |
| 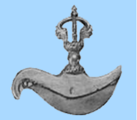 | 85 | altes Ritual, hacken [ancient ritual, chopping] | Spannung, erzeugen [electric current, making] |
| 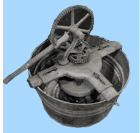 | 86 | Kleidung, im Eimer waschen [clothes, washing in a bucket] | Türgelenk, Feuer überstehen [door hinge, surviving fires] |
| 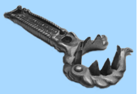 | 87 | Maiskolben, entkörnen [corncob, removing grains] | Brief, wiegen [letter, weighing] |
| 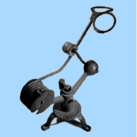 | 88 | Ball, katapultieren [ball, catapulting] | Radiergummi, mit Strom betreiben [eraser, operating with electricity] |
| 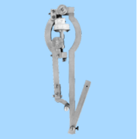 | 89 | Zeichnungen, vermessen [drawings, measuring] | Brenner, löten [burner, soldering] |

Table 1 continued

| Stimulus | ID | Matching keywords | Non-matching keywords |
|---|---|---|---|
| 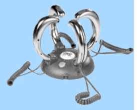 | 90 | Elektroschock, spielen [electric shock, playing] | Weinfass, Loch einschlagen [wine barrel, smashing a hole] |
| 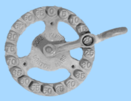 | 91 | Lieferungen, abzählen [deliveries, counting] | Außenbereich, heizen [outdoor area, heating] |
| 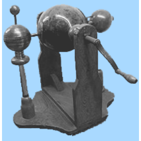 | 92 | Rotation, Ladung erzeugen [rotation, creating electric charge] | Musikgerät, stampfen [musical instrument, stamping] |
| 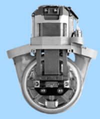 | 93 | Herz, durch Maschine ersetzen [heart, replacing with machine] | Kerzen, löschen [candles, extinguishing] |
| 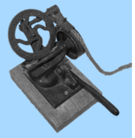 | 94 | Seil, schneiden [rope, cutting] | Akkordeon, spielen [accordion, playing] |
| 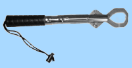 | 95 | Fisch, wiegen [fish, weighing] | Herz, durch Maschine ersetzen [heart, replacing with machine] |
| 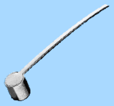 | 96 | Kerzen, löschen [candles, extinguishing] | Körper, trainieren [body, examining] |
| 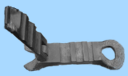 | 97 | Korken, formen [bottle cork, forming] | Sternenbilder, vermessen [stellar constellation, measuring] |
| 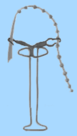 | 98 | Erektion, helfen [erection, helping] | Waffe, werfen [weapon, throwing] |
| 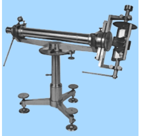 | 99 | Streckenmaß, Sonnenlicht nutzen [distance measure, using sunrays] | Kartoffeln, stampfen [potatoes, mashing] |
| 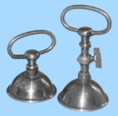 | 100 | Luftdruck, messen [air pressure, measuring] | Knochen, sägen [bones, sawing] |
| 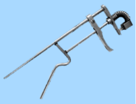 | 101 | Piano, stimmen [piano, tuning] | Eierkarton, pressen [egg carton, pressing] |
| 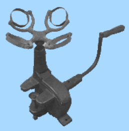 | 102 | von Hand, zentrifugieren [by hand, centrifugating] | Lieferungen, abzählen [deliveries, counting] |

Table 1 continued

| Stimulus | ID | Matching keywords | Non-matching keywords |
|---|---|---|---|
| 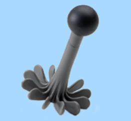 | 103 | Kartoffeln, stampfen [potatoes, mashing] | Nachricht, morsen [message, signaling] |
| 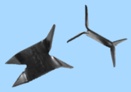 | 104 | Waffe, werfen [weapon, throwing] | elektrische Spannung, prüfen [electric current, measuring] |
| 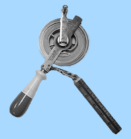 | 105 | Tankfüllstand, messen [fuel tank level, gauging] | Tonpott, trommeln [clay pot, drumming] |
| 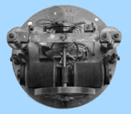 | 106 | Uhrzeit, anzeigen [time, displaying] | altes Ritual, hacken [ancient ritual, chopping] |
| 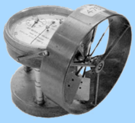 | 107 | Windgeschwindigkeit, messen [wind speed, measuring] | Waffe, entflammen [weapon, inflaming] |
| 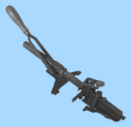 | 108 | Schlüsselloch, stanzen [keyhole, punching] | Automat, Eier ausbrüten [machine, incubating eggs] |
| 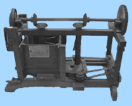 | 109 | Ziegelsteine, formen [bricks, forming] | Schallplatte, abtasten [vinyl record, reading] |
| 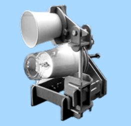 | 110 | Becher, Schall auffangen [drinking cup, picking up sound] | Schlüsselloch, stanzen [keyhole, punching] |
| 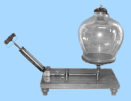 | 111 | Luft, abpumpen [air, pumping out] | Briketts, pressen [briquette, pressing] |
| 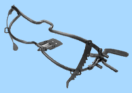 | 112 | Mund, offen halten [mouth, keeping open] | Orgelton, erzeugen [organ sound, making] |
| 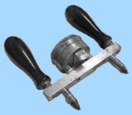 | 113 | Autobatterie, Spannung testen [car battery, measuring voltage] | Becher, Schall auffangen [drinking cup, picking up sound] |
| 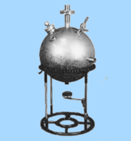 | 114 | Sonnenlicht, Intensität messen [sunlight, measure intensity] | Krawatten, aufhängen [necktie, hanging up] |
| 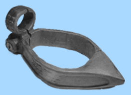 | 115 | Kutschrad, anschließen [carriage wheel, locking] | Angelschnur, kurbeln [fishing line, winding] |

Table 1 continued

| Stimulus | ID | Matching keywords | Non-matching keywords |
| --- | --- | --- | --- |
| 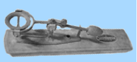 | 116 | Eier, wiegen<br>[eggs, weighing] | Leierkasten, klingen<br>[barrel organ, making sounds] |
| 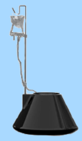 | 117 | Blumentopf, sich selbst wässern<br>[flowerpot, self-watering] | Glas, schneiden<br>[glass, cutting] |
| 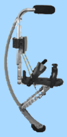 | 118 | Schritte, vergrößern<br>[steps, extending] | Tabletten, zerteilen<br>[pill, splitting] |
| 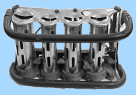 | 119 | Münzen, aufbewahren<br>[coins, storing] | Kutschrad, anschließen<br>[carriage wheel, locking] |
| 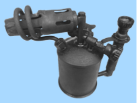 | 120 | Brenner, löten<br>[burner, soldering] | Flaschenkorken einführen, einführen<br>[bottle cork, inserting] |

### Online Results 1. *Linear Mixed-Effects Models*

#### *P1 Component (100–150 ms)*

```
## Linear mixed model fit by REML. t-tests use Satterthwaite's method [
## lmerModLmerTest]
## Formula: P1 ~ 1 + phase + condition + phase:condition + cosine + (1 +
##     phase | participant_id)
## Control: lme4::lmerControl
##
## REML criterion at convergence: 87306.7
##
## Scaled residuals:
##     Min      1Q  Median      3Q     Max
## -6.4625 -0.6237 -0.0031  0.6134  5.8089
##
## Random effects:
##  Groups         Name        Variance Std.Dev. Corr
##  participant_id (Intercept)  8.991   2.999
##                 phase1       1.670   1.292    -0.31
##                 phase2       1.822   1.350     0.44 -0.97
##  Residual                   25.582   5.058
## Number of obs: 14309, groups:  participant_id, 48
##
## Fixed effects:
##                     Estimate Std. Error        df t value Pr(>|t|)
## (Intercept)        3.991e+00  4.386e-01  4.857e+01   9.099 4.55e-12 ***
## phase1            -6.827e-01  2.156e-01  4.797e+01  -3.167  0.00268 **
## phase2             1.100e+00  2.230e-01  4.807e+01   4.930 1.02e-05 ***
## condition1         5.105e-02  1.054e-01  1.417e+04   0.484  0.62808
## condition2         1.790e-03  1.219e-01  1.418e+04   0.015  0.98829
## cosine             1.949e-01  7.413e-02  1.416e+04   2.629  0.00859 **
## phase1:condition1 -2.392e-01  2.553e-01  1.350e+04  -0.937  0.34874
## phase2:condition1  7.278e-01  2.563e-01  1.342e+04   2.840  0.00452 **
```

```
## phase1:condition2 -3.668e-01  2.933e-01  1.185e+04  -1.251  0.21113
## phase2:condition2  7.975e-01  2.946e-01  1.164e+04   2.707  0.00680 **
## ---
## Signif. codes:  0 '***' 0.001 '**' 0.01 '*' 0.05 '.' 0.1 ' ' 1
##
## Correlation of Fixed Effects:
##             (Intr) phase1 phase2 cndtn1 cndtn2 cosine phs1:1 phs2:1 phs1:2
## phase1      -0.265
## phase2       0.378 -0.857
## condition1   0.018  0.002  0.001
## condition2  -0.006  0.001  0.000  0.591
## cosine       0.127  0.000  0.001 -0.029 -0.042
## phs1:cndtn1  0.001  0.110 -0.053  0.002  0.002  0.002
## phs2:cndtn1  0.000 -0.055  0.110  0.006  0.005 -0.001 -0.511
## phs1:cndtn2  0.000 -0.002  0.002  0.002  0.002  0.000  0.582 -0.306
## phs2:cndtn2  0.000  0.002 -0.001  0.005  0.008  0.001 -0.306  0.584 -0.519
##
## Type III Analysis of Variance Table with Satterthwaite's method
##                 Sum Sq Mean Sq NumDF   DenDF F value    Pr(>F)
## phase           730.61  365.30     2    49.7 14.2799 1.258e-05 ***
## condition         8.91    4.46     2 14172.1  0.1742  0.840161
## cosine          176.75  176.75     1 14159.0  6.9091  0.008585 **
## phase:condition 259.50   64.88     4 11197.7  2.5360  0.038129 *
## ---
## Signif. codes:  0 '***' 0.001 '**' 0.01 '*' 0.05 '.' 0.1 ' ' 1
##
## Pairwise Contrasts (Simple Effects)
## contrast = Informed - Uninformed:
##  phase        estimate    SE    df t.ratio p.value
##  Pre-insight   -0.0321 0.180 12943  -0.178  1.0000
##  Insight       -0.2713 0.182 14138  -1.494  0.4060
##  Post-insight   0.4565 0.181 13047   2.515  0.0357
##
## contrast = Informed - Unsuccessful:
##  phase        estimate    SE    df t.ratio p.value
##  Pre-insight   -0.0195 0.206 10410  -0.095  1.0000
##  Insight       -0.3863 0.209 13860  -1.844  0.1956
##  Post-insight   0.4112 0.209 10673   1.972  0.1460
##
## Degrees-of-freedom method: satterthwaite
## P value adjustment: bonferroni method for 3 tests
```

#### *N170 Component (150–200 ms)*

```
## Linear mixed model fit by REML. t-tests use Satterthwaite's method [
## lmerModLmerTest]
## Formula: N170 ~ 1 + phase + condition + phase:condition + cosine + (1 +
##     phase | participant_id)
## Control: lme4::lmerControl
##
## REML criterion at convergence: 86644.7
##
## Scaled residuals:
##     Min      1Q  Median      3Q     Max
## -6.0470 -0.6299 -0.0035  0.6025  5.9459
##
## Random effects:
##  Groups         Name        Variance Std.Dev. Corr
##  participant_id (Intercept) 13.660   3.696
##                 phase1       6.588   2.567     0.15
##                 phase2       5.724   2.393     0.01 -0.96
##  Residual                   24.250   4.924
## Number of obs: 14309, groups:  participant_id, 48
##
```

```
## Fixed effects:
##                     Estimate Std. Error         df t value Pr(>|t|)
## (Intercept)        1.810e+00  5.379e-01  4.801e+01   3.365 0.001511 **
## phase1            -3.057e+00  3.851e-01  4.751e+01  -7.938 2.93e-10 ***
## phase2             2.611e+00  3.612e-01  4.706e+01   7.229 3.65e-09 ***
## condition1        -1.670e-01  1.026e-01  1.417e+04  -1.627 0.103705
## condition2        -2.168e-01  1.187e-01  1.417e+04  -1.826 0.067843 .
## cosine             3.197e-01  7.218e-02  1.416e+04   4.430 9.51e-06 ***
## phase1:condition1 -5.275e-01  2.497e-01  1.413e+04  -2.113 0.034637 *
## phase2:condition1  9.032e-01  2.505e-01  1.402e+04   3.606 0.000313 ***
## phase1:condition2 -8.385e-01  2.879e-01  1.384e+04  -2.913 0.003586 **
## phase2:condition2  1.198e+00  2.888e-01  1.341e+04   4.148 3.37e-05 ***
## ---
## Signif. codes:  0 '***' 0.001 '**' 0.01 '*' 0.05 '.' 0.1 ' ' 1
##
## Correlation of Fixed Effects:
##             (Intr) phase1 phase2 cndtn1 cndtn2 cosine phs1:1 phs2:1 phs1:2
## phase1       0.143
## phase2       0.006 -0.922
## condition1   0.014  0.001  0.001
## condition2  -0.005  0.001  0.000  0.592
## cosine       0.101  0.000  0.000 -0.029 -0.042
## phs1:cndtn1  0.000  0.059 -0.032  0.003  0.003  0.002
## phs2:cndtn1  0.000 -0.030  0.065  0.005  0.005  0.000 -0.508
## phs1:cndtn2  0.000 -0.002  0.001  0.003  0.003  0.000  0.587 -0.304
## phs2:cndtn2  0.000  0.001 -0.001  0.005  0.007  0.001 -0.304  0.588 -0.514
##
## Type III Analysis of Variance Table with Satterthwaite's method
##                  Sum Sq Mean Sq NumDF   DenDF F value    Pr(>F)
## phase           1529.29  764.64     2    48.1 31.5319 1.784e-09 ***
## condition         92.04   46.02     2 14166.2  1.8976 0.1499606
## cosine           475.81  475.81     1 14157.3 19.6209 9.514e-06 ***
## phase:condition  489.34  122.34     4 13093.5  5.0448 0.0004633 ***
## ---
## Signif. codes:  0 '***' 0.001 '**' 0.01 '*' 0.05 '.' 0.1 ' ' 1
##
## Pairwise Contrasts (Simple Effects)
## contrast = Informed - Uninformed:
##  phase        estimate    SE    df t.ratio p.value
##  Pre-insight   -0.1163 0.176 13762  -0.662  1.0000
##  Insight       -0.6439 0.177 14194  -3.631  0.0008
##  Post-insight   0.2593 0.177 13742   1.462  0.4312
##
## contrast = Informed - Unsuccessful:
##  phase        estimate    SE    df t.ratio p.value
##  Pre-insight   -0.0571 0.202 12610  -0.282  1.0000
##  Insight       -0.8956 0.205 14203  -4.371  <.0001
##  Post-insight   0.3024 0.204 12556   1.479  0.4171
##
## Degrees-of-freedom method: satterthwaite
## P value adjustment: bonferroni method for 3 tests
```

#### *N400 Component (400–700 ms)*

```
## Linear mixed model fit by REML. t-tests use Satterthwaite's method [
## lmerModLmerTest]
## Formula: N400 ~ 1 + phase + condition + phase:condition + cosine + (1 +
##     phase | participant_id)
## Control: lme4::lmerControl
##
## REML criterion at convergence: 82101.6
##
## Scaled residuals:
##     Min      1Q  Median      3Q     Max
```

```
## -6.7184 -0.6289 -0.0010  0.6330  5.0726
##
## Random effects:
##  Groups         Name        Variance Std.Dev. Corr
##  participant_id (Intercept)  2.842   1.686
##                 phase1       1.566   1.251     0.19
##                 phase2       1.013   1.007    -0.03 -0.61
##  Residual                   17.754   4.214
## Number of obs: 14309, groups:  participant_id, 48
##
## Fixed effects:
##                     Estimate Std. Error         df t value Pr(>|t|)
## (Intercept)        1.145e+00  2.504e-01  5.053e+01   4.571 3.16e-05 ***
## phase1             2.628e+00  2.019e-01  4.746e+01  13.018  < 2e-16 ***
## phase2            -1.615e+00  1.712e-01  4.842e+01  -9.433 1.51e-12 ***
## condition1         5.837e-01  8.777e-02  1.418e+04   6.651 3.02e-11 ***
## condition2         5.025e-01  1.015e-01  1.420e+04   4.950 7.51e-07 ***
## cosine             5.312e-02  6.176e-02  1.416e+04   0.860 0.389735
## phase1:condition1  1.216e+00  2.135e-01  1.407e+04   5.694 1.27e-08 ***
## phase2:condition1 -2.903e-01  2.140e-01  1.383e+04  -1.356 0.175088
## phase1:condition2  1.317e+00  2.460e-01  1.362e+04   5.353 8.80e-08 ***
## phase2:condition2 -8.411e-01  2.465e-01  1.286e+04  -3.412 0.000647 ***
## ---
## Signif. codes:  0 '***' 0.001 '**' 0.01 '*' 0.05 '.' 0.1 ' ' 1
##
## Correlation of Fixed Effects:
##             (Intr) phase1 phase2 cndtn1 cndtn2 cosine phs1:1 phs2:1 phs1:2
## phase1       0.162
## phase2      -0.027 -0.585
## condition1   0.027  0.002  0.001
## condition2  -0.009  0.001  0.000  0.591
## cosine       0.185  0.000  0.001 -0.029 -0.042
## phs1:cndtn1  0.001  0.097 -0.058  0.003  0.003  0.002
## phs2:cndtn1  0.000 -0.049  0.119  0.005  0.004  0.000 -0.500
## phs1:cndtn2  0.000 -0.003  0.001  0.003  0.003  0.000  0.586 -0.294
## phs2:cndtn2  0.000  0.001 -0.001  0.004  0.006  0.001 -0.294  0.587 -0.501
##
## Type III Analysis of Variance Table with Satterthwaite's method
##                 Sum Sq Mean Sq NumDF   DenDF F value    Pr(>F)
## phase          3097.77 1548.88     2    48.1 87.2427 < 2.2e-16 ***
## condition       813.58  406.79     2 14180.1 22.9129 1.162e-10 ***
## cosine           13.13   13.13     1 14159.6  0.7398    0.3897
## phase:condition  831.16  207.79     4 13624.4 11.7040 1.728e-09 ***
## ---
## Signif. codes:  0 '***' 0.001 '**' 0.01 '*' 0.05 '.' 0.1 ' ' 1
##
## Pairwise Contrasts (Simple Effects)
## contrast = Informed - Uninformed:
##  phase        estimate    SE    df t.ratio p.value
##  Pre-insight    -0.130 0.151 14117  -0.861  1.0000
##  Insight         1.086 0.151 14156   7.173  <.0001
##  Post-insight    0.795 0.152 14031   5.230  <.0001
##
## contrast = Informed - Unsuccessful:
##  phase        estimate    SE    df t.ratio p.value
##  Pre-insight    -0.095 0.174 13794  -0.546  1.0000
##  Insight         1.222 0.175 13925   6.998  <.0001
##  Post-insight    0.381 0.176 13495   2.169  0.0903
##
## Degrees-of-freedom method: satterthwaite
## P value adjustment: bonferroni method for 3 tests
```

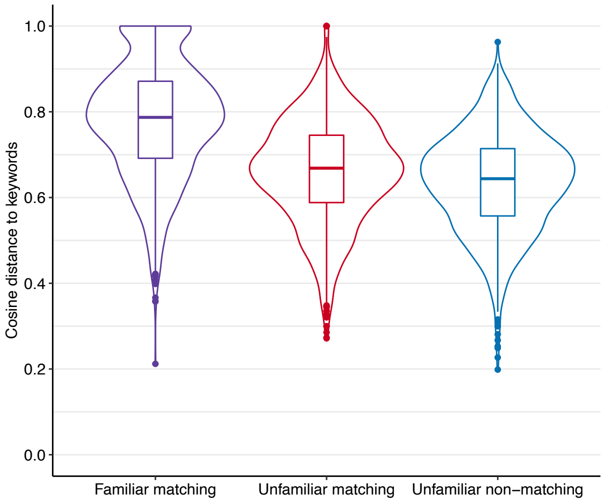


***Online Figure 1.*** Online pre-rating study results. Participants were presented with 240 objects in random order, 120 of which were familiar everyday objects and the other 120 were presumed to be unfamiliar to most people. Participants were asked to describe or guess the function of each object by typing a pair of German keywords. Violins show the distributions of the similarities between these participant-generated keywords and the keywords that we had created for our main EEG experiment (see Materials and Methods; Online Table 1). We computed these similarities separately for (a) participant-generated keywords for the familiar objects and keywords that we had created to match the familiar objects (though these were not part of the main EEG experiment; red), (b) participant-generated keywords for the unfamiliar objects and keywords that we had created to match the unfamiliar objects (blue), and (c) participant-generated keywords for the unfamiliar objects and keywords that we had created to not match the unfamiliar objects (by selecting keywords that matched one of the other unfamiliar objects; purple). Semantic similarities were computed as the cosine similarity between the sums of the two word vectors in a word2vec emebedding space pre-trained on the German Wikipedia. Boxplots show the median (thick line), 25th and 75th percentiles (hinges), 1.5 times the interquartile range above and below the hinges (whiskers), and any outlier data points that fall outside of the whiskers (dots).

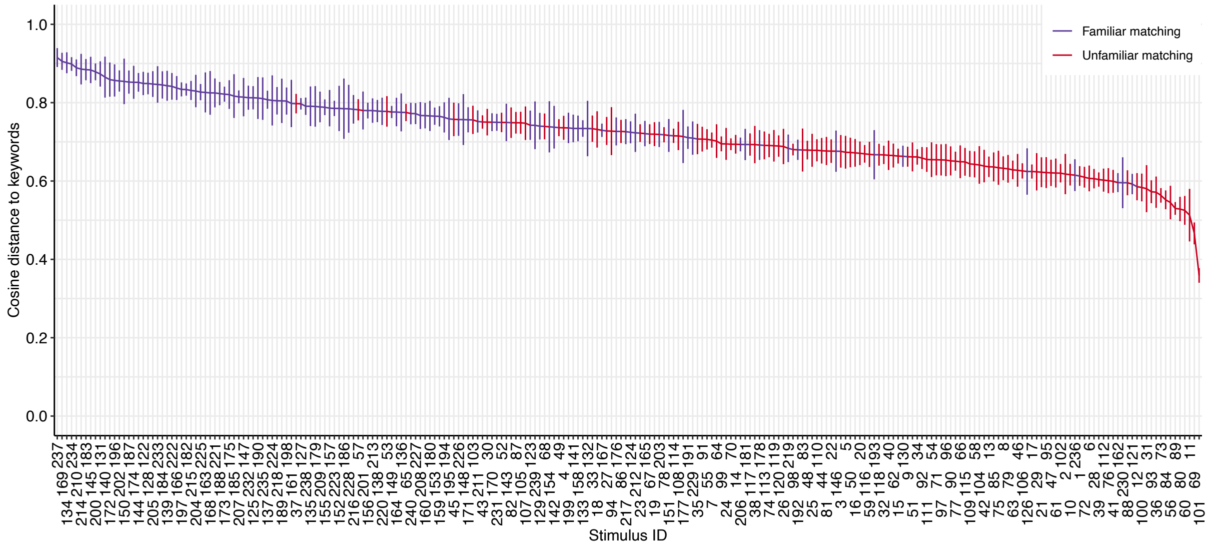


***Online Figure 2.*** Online pre-rating study results on the level of individual object stimuli. Same as Online Figure 1, but showing, for each object stimulus separately, the mean (horizontal line) ± 1 standard error (vertical lines) of the semantic similarities between participant-generated keywords and the keywords that we had generated. Semantic similarities were generally higher for the familiar objects than for the unfamiliar objects, indicating that it was easier for participants to come up with the correct function for the familiar objects. This mean cosine similarity for each object (i.e., a measure of the difficulty of guessing its function) was entered as a covariate of no interest in all linear mixed-effects models for analyzing the data of the main EEG experiment (see Materials and Methods; Results).

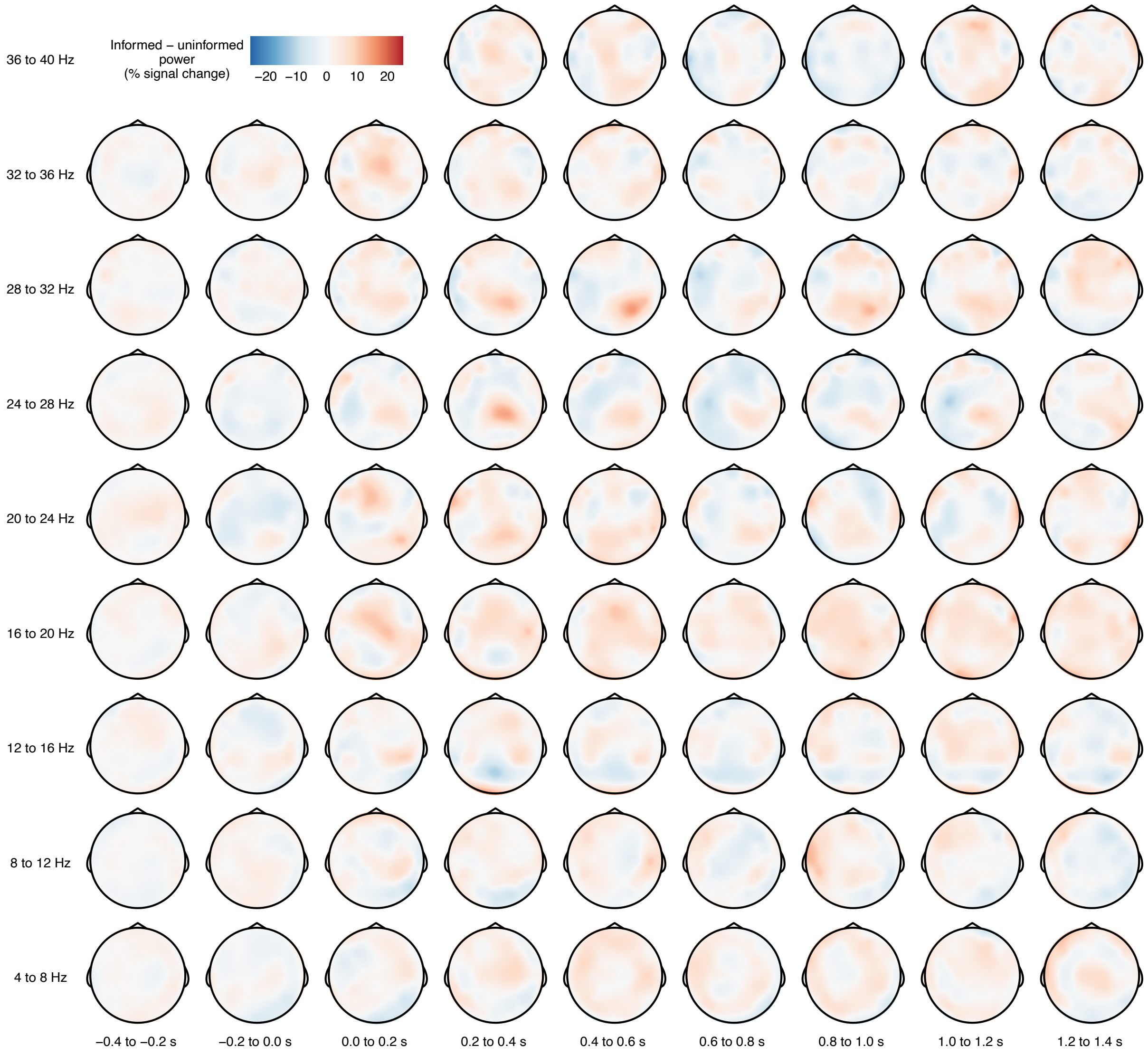


***Online Figure 3.*** Time-frequency results for the pre-insight phase. Each topographic plot shows the difference in event-related power (in units of percent signal change) between the semantically informed condition and the uninformed condition, grand-averaged across participants. A cluster-based permutation test indicated no clusters for which this difference was statistically significant (all $ps > .789$).

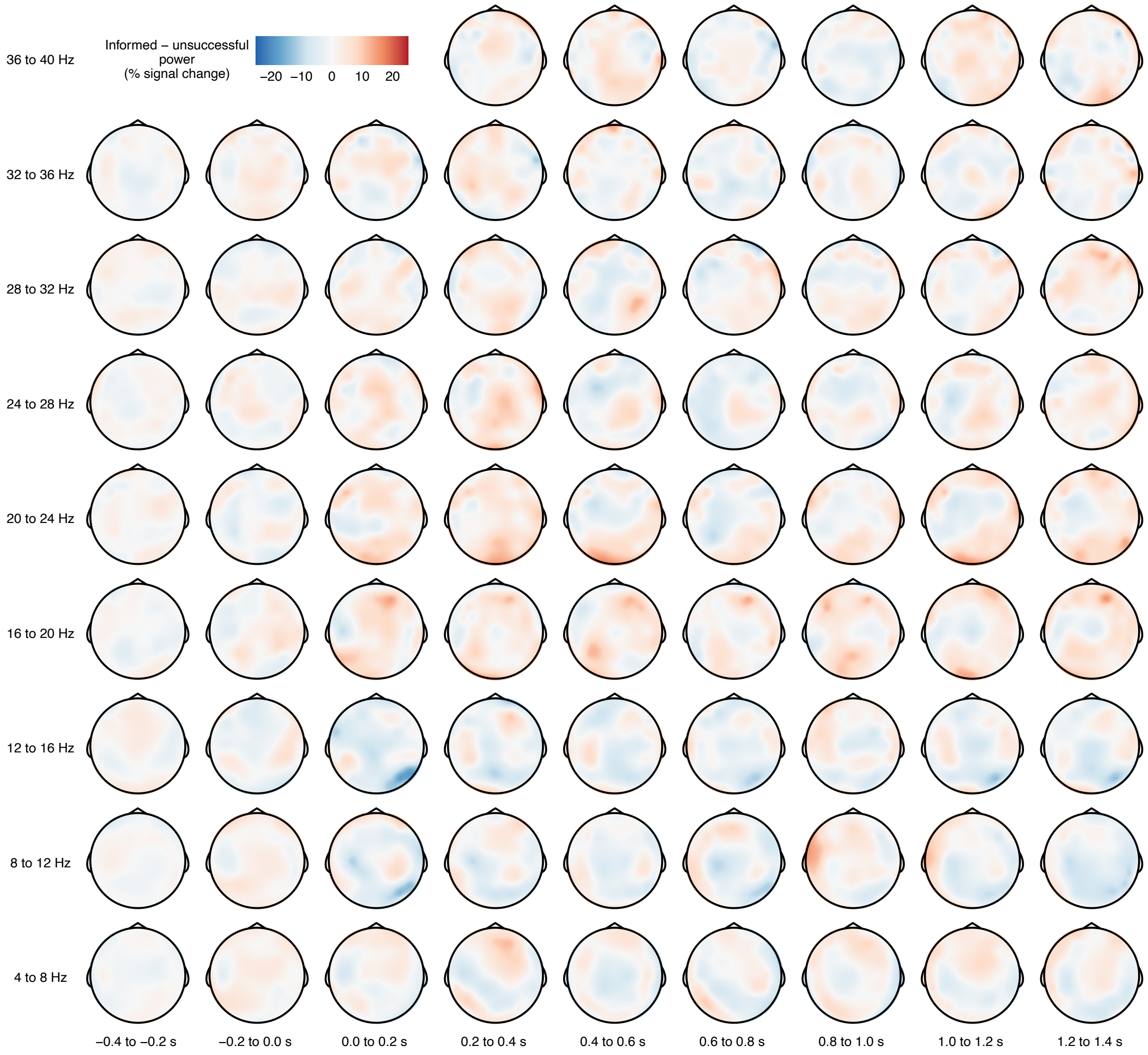


***Online Figure 4.*** Time-frequency results for the pre-insight phase (informed minus unsuccessful). Each topographic plot shows the difference in event-related power (in units of percent signal change) between the semantically informed condition and the unsuccessfully informed condition, grand-averaged across participants. A cluster-based permutation test indicated no clusters for which this difference was statistically significant (all *ps* $>$ .770).

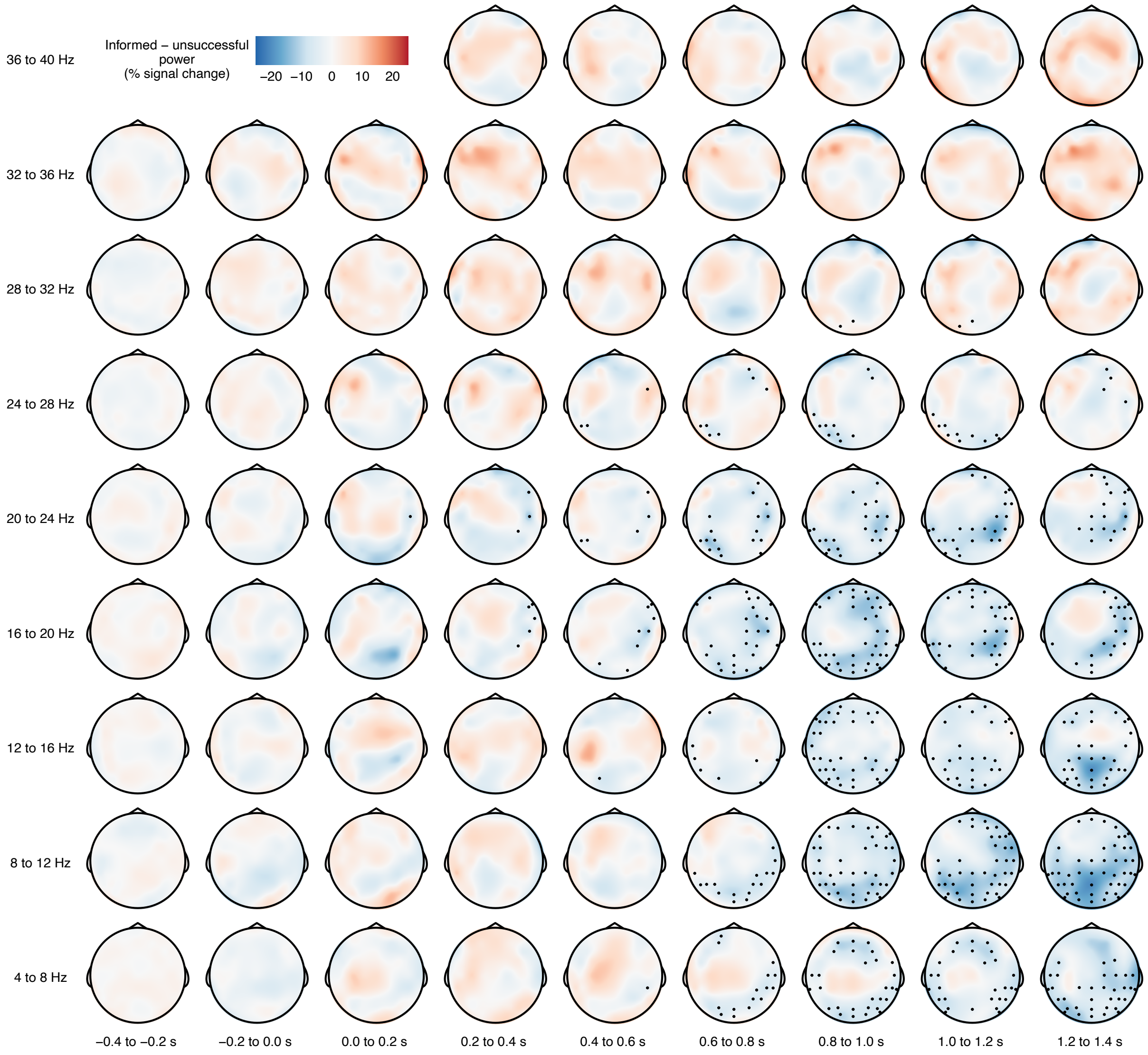


***Online Figure 5.*** Time-frequency results for the insight phase (informed minus unsuccessful). Each topographic plot shows the difference in event-related power (in units of percent signal change) between the semantically informed condition and the unsuccessfully informed condition, grand-averaged across participants. Black dots highlight EEG channels that were part of a cluster for which this difference was statistically significant ($p_{\text{cluster}}$ = .016).

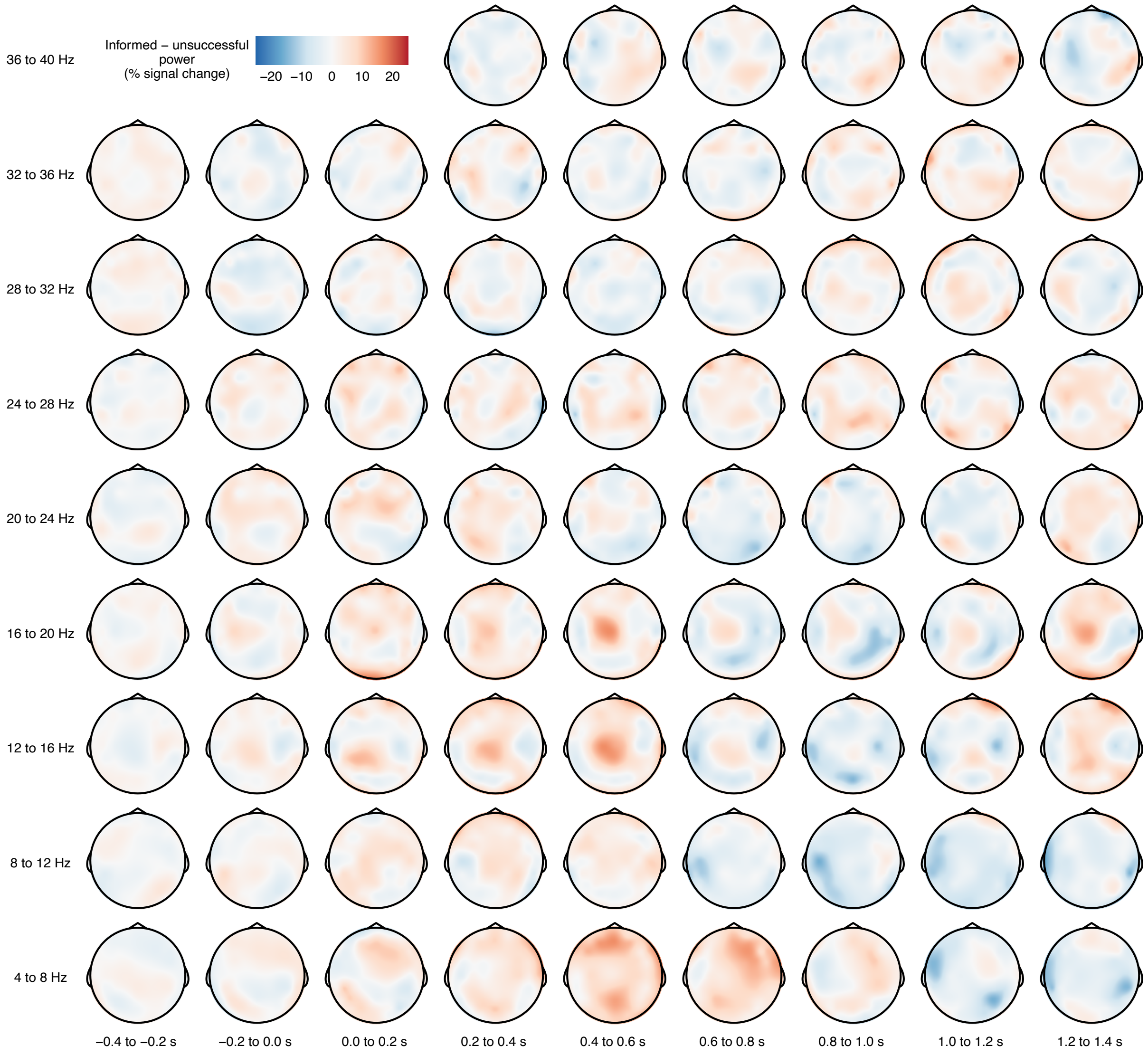


***Online Figure 6.*** Time-frequency results for the post-insight phase (informed minus unsuccessful). Each topographic plot shows the difference in event-related power (in units of percent signal change) between the semantically informed condition and the unsuccessfully informed condition, grand-averaged across participants. A cluster-based permutation test indicated no clusters for which this difference was statistically significant (all $ps > .216$).
